# Source tracking of larval bacterial community of Pacific white shrimp across the developmental cycle: ecological insights for microbial management and pathogen prevention in larviculture

**DOI:** 10.1101/2025.05.25.655974

**Authors:** Fengdie Zhang, Heng Chen, Junqi Yu, Rudan Chen, Demin Zhang, Chen Chen, Kai Wang

**Author notes:** For correspondence. E-mail Kai Wang or Chen Chen.

## Abstract

Health of postlarvae (as products of larviculture) of Pacific white shrimp (*Penaeus vannamei*) is critical to subsequent culture. Larval microbiota establishment has potential long-term impact on host health, but bacterial sources of larval shrimps remain largely unexplored. We investigated the larviculture cycles of two shrimp strains by utilizing 16S rRNA gene sequencing alongside mainstream microbial source-tracking tools. Our goal was to evaluate the contributions of environmental elements, feeds, and internal succession to larval bacterial community. Regardless of shrimp strains or tools, the mouth opening marked a pivotal shift in bacterial source composition of larvae, with a notable contribution from the rearing water at the beginning of mouth opening, followed by enhanced internal succession, emphasizing the critical role of early microbiota establishment. Notably, the dominance of external sources at *zoea* II highlights the necessity for ensuring microbial safety due to the prevalence of *zoea*-II syndrome at this vital stage. Source tracking of specific bacterial groups suggests regulatory timing and routes of *Roseobacteraceae* (with potential probiotics) and frozen *Artemia* as a major source of vibrios (with opportunistic pathogens). The determinism-dominated assembly of bacteria, sourced either internally or externally, highlights strong host selection. This process exhibits overall similarities in taxonomic preferences for internal succession, while differences in taxonomic preferences from external sources reflect the influence of the genetic backgrounds of shrimp strains. Our work presents the first systematic assessment on bacterial sources of *P. vannamei* larvae across the complete developmental cycle, providing new insights that facilitate microbial management and pathogen prevention in larviculture.

## 1. Introduction

The culture of Pacific white shrimp (*Penaeus vannamei*, formerly *Litopenaeus vannamei*), as the most productive prawn species in world aquaculture industry (FAO, 2024), has been consistently suffered from frequent disease outbreaks (Kumar et al., 2020; Sha et al., 2022) and decline of stress resistance (Duan et al., 2024) for the recent decade. Empirically and practically, the health and quality of postlarval shrimps (as indicated by survival rate) after a complete larviculture cycle (around 15∼16 days, covering the developmental stages including *nauplius*, *zoea*, *mysis*, and early *postlarvae*) is closely related to the growth performance and stress resistance of shrimps in subsequent culture, thus likely determining the success of shrimp culture to large extent (Racotta et al., 2004; Racotta et al., 2003). It is generally believed that the health and quality of postlarval shrimps is simultaneously determined by genetic background and environmental regulation during larviculture (Emerenciano et al., 2022).

Microbial community serve as a crucial hub between the environment of culture system and aquaculture animals (El-Saadony et al., 2022; Holt et al., 2021). A growing number of studies have demonstrated the associations and/or causality of microbial communities of gut and/or rearing water with health status (He et al., 2020; Huang et al., 2018), growth and development (Fan et al., 2019), and disease/stress resistance (Guo et al., 2023) of *P. vannamei*. Some studies have found that microbial community disorders in early life across a spectrum of animals, including human (infants and young children) (Backhed et al., 2015), mice (Hughes et al., 2020), calves (Ma et al., 2020), and oysters (Dai et al., 2023), have potential long-term impact on host health. For example, the impact of early life events (like antibiotics exposure) to gut microbiome of human infants has been reported to be associated with obesity (Kerr et al., 2015), chronic enteritis (Kronman et al., 2012), and immune diseases (Healy et al., 2022) in later life stages. The establishment of microbiota of shrimp larvae may also have potential long-term impact on their health in subsequent culture (Zhang et al., 2021). Therefore, microbial management should be considered as an important environmental regulatory strategy (and eventually as a routine) for achieving beneficial larval microbiota in larviculture practice. After naupliar shrimps are hatched, the microbial communities in the gut, epidermis, and other tissues and organs are gradually assembled and then established. Identifying microbial sources of larval shrimps and timing and processes of their entry into the host during the complete developmental cycle can facilitate clarification of regulatory pathway and timing for larval shrimp microbiota, and thus is essential for the practice of microbial management.

Knowing sources of microbiota (especially health-related microbial signature taxa including potential probiotics and pathogens) of larval shrimps is not only a key to reveal mechanisms underlying larval microbial community assembly, but is also fundamental to understand who, when, and how the host tend to select for or against during the rapid development, thus providing guidelines for beneficial microbial management and pathogen prevention. A previous study found that bacterial community assembly in the rearing water of *P. vannamei* larvae was dominated by stochastic processes, and 37% of bacterioplankton were introduced by live and dry feeds and exchange water (Heyse et al., 2021). While, approximately 66.7% of prokaryotes in the gut of adult shrimps were succeeded from juvenile shrimps as inferred by SourceTracker (Zhang et al., 2021). Our previous study further indicated the impact of rearing water (especially the water from the previous stage) on bacterial communities of larval shrimps and positive host selection for certain bacteria (such as *Rhodobacteraceae* taxa) from the rearing water at certain stages (especially *zoea* sub-stages) (Wang, Y. et al., 2020). Also, the assembly of larval bacterial community was overall governed by neutral processes (dispersal among individuals and ecological drift) when considering the larval bacterial metacommunity at a given stage as the source community (Wang, Y. et al., 2020). These findings indicate certain stochasticity in bacterial community assembly in the larviculture system (both water and larvae) may be related to health status differentiation of larval shrimps even with uniform managements in practice, enlightening us to realize that the goal of microbial regulation during the laviculture period is to strengthen deterministic assembly processes of microbial community. To achieve this purpose, we must firstly find out which microbial sources and groups colonize larval shrimps through which ecological processes. However, the relative contribution of potential microbial sources, involving environmental elements and inputs in(to) the larviculture system as well as internal succession from larvae of previous developmental stages to the larval microbial community across the complete developmental cycle has not been comprehensively evaluated.

In this study, we monitored the complete larviculture cycle of two strains of *P. vannamei* with different culture traits (SR: stress-resistant strain; FG: fast-growing strain). Larval shrimps at different developmental stages and potential sources of larval microbiota including environmental elements and inputs in(to) the larviculture system were comprehensively collected. Based on 16S rRNA gene amplicon sequencing and two mainstream tools for microbial source tracking, we aimed (1) to assess the relative contribution of potential sources at multiple category levels to the bacterial community and major bacterial groups of larval shrimps; (2) to estimate ecological processes and timing by which and when bacteria were assembled from potential sources into the larval bacterial community; and (3) to identify commonness and difference in taxonomic selection from bacterial sources by two shrimp strains.

## 2. Materials and methods

### 2.1 Experimental design, sample collection, and monitoring of water environmental conditions

The management of the laviculture cycle and the sampling of shrimp larvae and rearing waters have been described in our recent work (Chen et al., 2025), albeit with different focuses. The fertilized eggs of two strains of *P. vannamei* (SR: stress-resistant strain; FG: fast-growing strain) were spawned and then hatched in Qingjiang Aquaculture Base, Zhejiang Mariculture Research Institute, Wenzhou, China (28.28°N, 121.11°E). During the selective breeding process, the SR strain was selected under low-salinity stress (∼3 psu) based on breeding values for survival rate, serving as an indicator of stress resistance, while the FG strain was selected based on breeding values for growth performance. After hatching, the naupliar shrimps (*nauplius* I) were immediately transferred to Yongxing Aquaculture Base, Zhejiang Mariculture Research Institute (27.859°N, 120.845°E), for larviculture. Six standardized ponds (7 m × 5 m × 2 m) in a larviculture workshop were randomly divided into two groups for the two shrimp strains, as three biological replicates for each strain (two replicates remaining for FG strain after 10 days post hatching (dph) due to the extinction of larvae in one of the FG ponds by 10 dph as described below). The initial density of naupliar shrimps was approximately 1.43×10^5^ nauplii/m^3^.

During the period of larviculture, lasting for 15 dph, the rearing water was constantly aerated and maintained under the following conditions: temperature 26.3-33.1 °C, pH 8.0-8.4, dissolved oxygen 6.4-7.4 mg/L, and salinity 15.1-28.8 psu. Water temperature and salinity were turned down to approximately 26 °C and 15 psu at *postlarvae* V for customized commercial purpose of preadaptation to the subsequent culture. The dynamics of inorganic nitrogen in the rearing water were characterized in another study of ours investigating the divergence patterns of the larval bacterial community (Chen et al., 2025), with similar initial levels—ammonia nitrogen at 0.554 ± 0.266 (mean ± standard error) and 0.321 ± 0.068 mg/L, nitrite nitrogen at 0.005 ± 0.003 and 0.021 ± 0.004 mg/L, and nitrate nitrogen at 0.224 ± 0.022 and 0.190 ± 0.040 mg/L— in SR and FG ponds. The divergence in inorganic nitrogen levels between the ponds culturing the two shrimp strains during the later stages—specifically, the higher levels of ammonia from *mysis* I and the higher levels of nitrite and nitrate from *postlarvae* I in FG ponds compared to SR ponds—may be partially attributed to differences in the feeding behavior of the two strains, which in turn lead to varying amounts of feed residue. However, as feeding behavior was not monitored in this study, this remains a speculative explanation.

All the ponds were maintained by the uniform management including the regulation of environmental conditions, inputs, and water exchange. The feeding and water-exchange regimes during the larviculture are shown in Fig.1. Specifically, the larvae were fed with live microalgae, including *Chaetoceros* sp., *Isochrysis* sp., and *Dirateria* sp., at a density of approximately 10^10^ cells/m^3^ from the transitional stage of *nauplius* VI - *zoea* I to *zoea* I stage (1 time/day). Commercial shrimp flakes, mixed feed, polysaccharide, vitamin C powder, and spirulina powder were applied after mouth opening at *zoea* I (3 times/day), following the frequency shown in Fig.1. The larvae were also fed with frozen *Artemia* from *zoea* II to *mysis* (11 dph) and with live *Artemia* from *postlarvae* I to *postlarvae* (14 dph). We obtained *Artemia* eggs from a commercial supplier and hatched them in-house. Frozen and live *Artemia* were applied at a density of approximately 2.86×10^5^ individuals and 5.72×10^5^ individuals/m^3^, respectively (both 3 times/day). Exchange water (∼1,750 L) was introduced to each pond at *mysis* I, *postlarvae* I, and *postlarvae* (13 dph).

**Figure 1.**
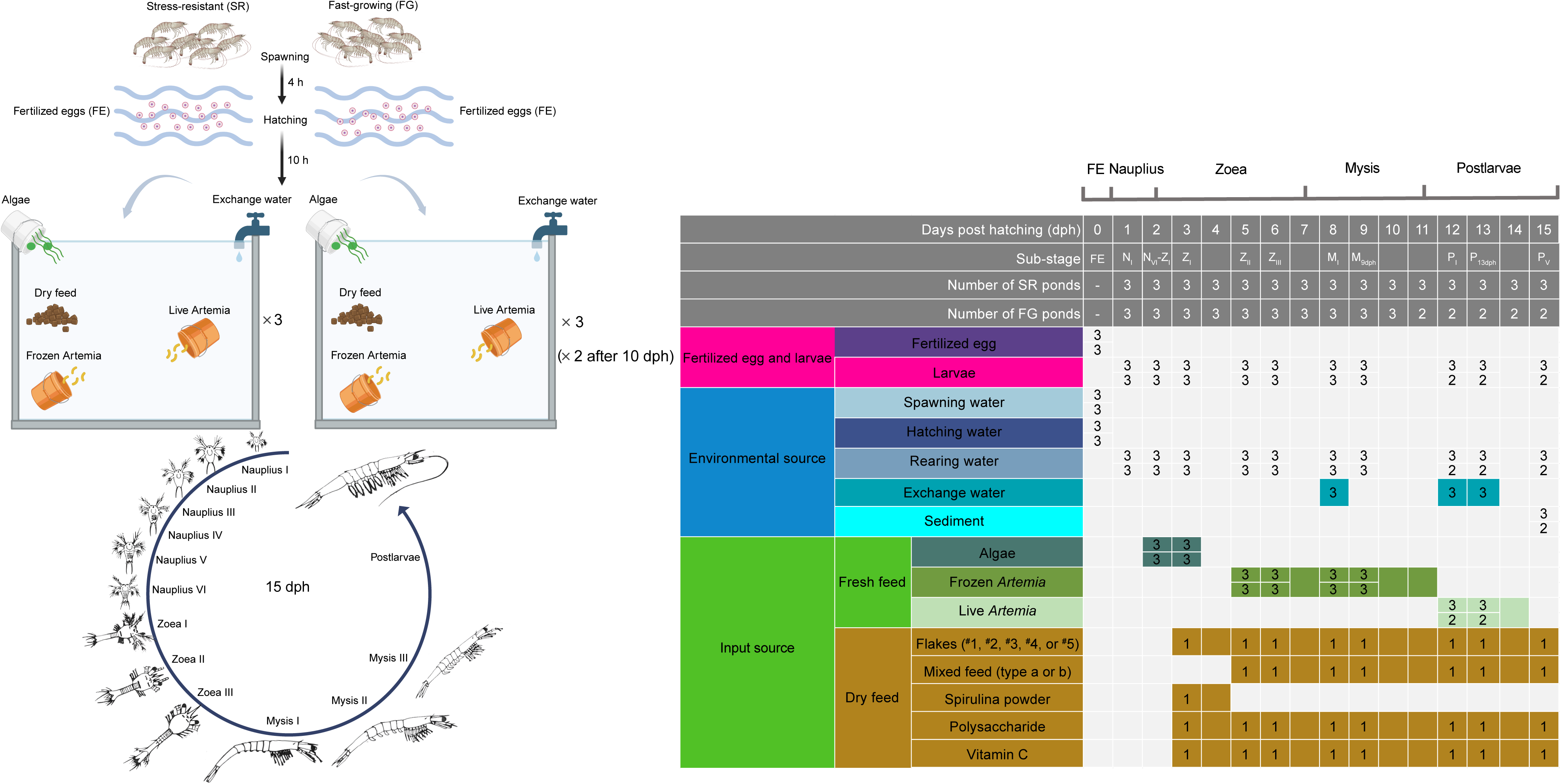
The sampling scheme across the complete developmental cycle of shrimp larvae. Shrimp larvae of both strains across different developmental stages (including fertilized egg (FE), *nauplius* (N), *zoea* (Z), *mysis* (M), and *postlarvae* (P)) and the potential sources (including environmental and input sources) expected to contribute to the larvae bacterial community were sampled at the days indicated by the cells filled with numbers, which represent the numbers of biological samples collected. In cells divided into upper and lower sections, the upper number denotes the number of biological replicates collected for the SR group, while the lower number corresponds to the FG group. Shaded cells represent the feeding and water-exchange regimes. N_VI_-Z_I_, transitional stage between *nauplius* VI and *zoea* I; M_9dph_, larvae samples from 9 dph (days post hatching) including mixed *mysis* sub-stages; P_13dph_, larvae samples from 13 dph including mixed *postlarvae* sub-stages.

Larval developmental stages were determined using microscopy. Specifically, 500 mL of water was collected from the water column of each pond, and larvae in the samples were identified based on morphological characteristics. If more than 80% of the individuals had completed metamorphosis to a specific stage, the sample was classified as being at that developmental stage; otherwise, it was considered to represent mixed stages (Wang, R. et al., 2020). Larvae from both shrimp strains were sampled throughout the entire developmental cycle—including fertilized eggs, *nauplius*, *zoea*, *mysis*, and early *postlarvae*—along with potential sources expected to contribute to the larval bacterial community, including environmental elements and feed inputs. The sampling strategy is illustrated in Fig. 1, where the numbers indicate the biological replicates collected for each sample type at different developmental stages. In cells divided into upper and lower sections, the upper number denotes the number of biological replicates collected for the SR group, while the lower number corresponds to the FG group. Sampled larvae (∼ 0.8 g fresh weight per sample [pond]) were gently washed with sterilized ultrapure water to remove the adsorbed rearing water, and then were transferred into sterilized 2-mL tubes. The *Artemia* samples were treated in the same way. 1 L of spawning, hatching, exchange, and rearing water samples were collected and prefiltered through a 160-mesh (∼95-μm) sterile nylon screen. The samples were then filtered onto a 0.2-μm polycarbonate membrane (Millipore, USA). The final volume of filtered water ranged from 0.2 to 1 L, depending on the turbidity of the water. Algal samples were collected before being poured into the ponds and then were filtered onto the 0.2-μm polycarbonate membrane. All the samples were collected and stored at −80 °C until DNA extraction. A total of 6 fertilized egg samples, 57 larvae samples, 83 environmental samples (including 6 spawning water samples, 6 hatching water samples, 57 rearing water samples, 9 exchange water samples, and 5 sediment samples), and 46 samples of fresh feeds (including 12 algal samples, 24 frozen *Artemia* samples, and 10 live *Artemia* samples) were used for DNA extraction and sequencing (Fig. 1). Although one biological sample of each type of dry feed was collected at each sampling stage during feeding, a single representative sample—sourced from the same package as all others of its type—was used for DNA extraction and sequencing, and served as the reference across all sampling stages for comparative purposes. In total, 10 dry feed samples were analyzed, including 5 flake samples (^#^1–^#^5), 2 mixed feed samples (types a and b), 1 spirulina powder sample, 1 polysaccharide sample, and 1 vitamin C sample. In summary, additional microbiome sequencing data from potential sources— including feed inputs and environmental elements beyond the rearing water—were integrated with the larval and rearing water datasets previously reported by Chen et al. (2025).

At the end of larviculture (*postlarvae* V), the survival rate of larvae in each pond was recorded. Based on extensive experience in long-term larviculture practices at the base we conducted this study and previous studies within larviculture systems under farming conditions, the survival rates of healthy or symptom-free *Penaeus vannamei* larvae by the end of the full larviculture cycle (typically at the *postlarvae* V stage) generally range from 13.8% to 50.6% (Reyes et al., 2022; Wang, R. et al., 2020).

Other shrimp species, such as *Penaeus stylirostris*, displayed survival rates of less than 45% in most ponds, even when subjected to antibiotic treatments for just 9 dph (Callac et al., 2022). In this study, the final larval survival rates of the SR strain in all three ponds fell within the normal range (47.7%, 32.4%, and 31.0%). Similarly, the survival rates in two of the FG ponds were also within the expected range (29.6% and 27.0%). However, in one of the FG ponds, the larvae were nearly extinct by 10 dph (and the pond was dismissed to cut costs), with a recorded survival rate of 0%.

Despite this, no signs of disease were observed in this pond or any of the others. The overall lower survival rates of FG strain compared with SR strain (SR vs. FG: 37.1% vs. 18.9% in average) corresponded to the variation in their breeding values for survival rate (as an indicator of stress resistance; data not shown) (Chen et al., 2025). Although all the larval samples collected were maintained under normal conditions, a reduction in the number of biological replicates for the FG strain after 10 dph (during the *postlarvae* stage) may slightly weaken statistical power. However, the overall data trends across developmental stages are likely to remain robust. Despite a balance between statistical power and workload feasibility must be considered in real-world larviculture settings, future studies with a larger number of replicates would help enhance statistical power and lead to more robust conclusions.

### 2.2 DNA extraction, 16S rRNA gene amplification, and Illumina sequencing

Total DNA of fertilized eggs and larvae, frozen and live *Artemia*, and all the dry feeds was extracted using a QIAamp® DNA Stool Mini Kit (QIAGEN, Germany). Total DNA on the filters of rearing water, exchange water, and algal feed or in sediments was extracted using a SPINeasy DNA Kit for Soil (MPBIO, USA). The V4 region of 16S rRNA genes was amplified using the primers set 515FY (5’-GTGYCAGCMGCCGCGGTAA-3’) and 806RB (5’-GGACTACNVGGGTWTCTAAT-3’) with dual barcodes (Apprill et al., 2015; Parada et al., 2016). An amount of 50 ng purified DNA template from each sample was amplified with a 50-µl reaction system under the following conditions: initial denaturation at 95 °C for 3 min, then 28 cycles of denaturation at 95 °C for 30 s, annealing at 55 °C for 30 s, and extension at 72 °C for 45 s; with a final extension at 72 °C for 10 min. The PCR products were purified with an E.Z.N.A.® Gel Extraction Kit (Omega, USA). The library was prepared with an ALFA-SEQ DNA Library Prep Kit (FINDROP, China), quantified using a Qubit 4.0 fluorometer (Invitrogen, USA), and then sequenced on an Illumina NovaSeq 6000 platform (Illumina, USA) at MAGIGENE Co., Ltd., Guangzhou, China.

### 2.3 Sequence processing

Sequences were processed using USEARCH v11.0.667 as previously described (Yan et al., 2024). Briefly, the paired reads were joined using the script-*fastq_mergepairs*. The joined sequences were quality-checked (maximum expected errors =1.0) and de-replicated using the script *usearch11-fastx_uniques*. Then, denoise, chimera check and filter, and identification of Zero-radius Operational Taxonomic Units (ZOTUs, also known as Amplicon Sequence Variants (ASV)) were conducted using the script *usearch11-unoise3* (minsize =4) based on UNOISE3 algorithm(Edgar, 2016). The original joined reads were then mapped to the ZOTU sequences at 100% similarity using the script *usearch11-otutab* to obtain ZOTU abundances. Subsequently, the sequences were aligned using the script *-super5* based on MUSCLE5 (Edgar, 2022), and a phylogenetic tree was generated using FastTree (Price et al., 2010). The taxonomy of ZOTUs were assigned against the NCBI RefSeq ribosomal RNA loci database (download on April 20, 2024) (O’Leary et al., 2016) using Blastn v2.13.0+ (with a percentage of identity >75%, max_target_seqs =1, and an *e*-value < 0.0001). To target the bacterial community, Archaea, Mitochondria, Chloroplast, and unassigned sequences were removed from the ZOTU table. The full dataset (n = 211) after above procedures contained 13,597 ZOTUs with 17,600,102 qualified reads (ranging from 10,991 to 157,437 reads per sample except a sample with low read depth, mean 83,412 reads per sample). One of the algal samples at *zoea* I was discarded due to low sequencing depth (6,879 reads). The ZOTU table was rarefied at 10,900 reads per sample for downstream analyses unless otherwise stated.

### 2.4 Bacterial Source tracking of larval and water bacterial communities

Bacterial source tracking for shrimp larvae was conducted using both SourceTracker (Knights et al., 2011) and Fast Expectation-maximization Microbial Source Tracking (FEAST) (Shenhav et al., 2019) algorithms with the R packages ‘SourceTracker’ and ‘FEAST’ with default settings, to complementarily test the robustness of the results and to strengthen the solidness of conclusions. As two of the most popular microbial source tracking tools, SourceTracker is a Bayesian method that allows for uncertainty in the distribution of sources and sinks, explicitly models the sink as a convex mixture of sources and deduces the mixture proportions from Gibbs sampling (Knights et al., 2011). While, FEAST is a highly efficient expectation-maximization-based method that is more scalable than the Markov Chain Monte Carlo used by SourceTracker, specially has a more sensitive source allocation capability that can be used for precise control of sources and sinks (Shenhav et al., 2019). According to a comprehensive study comparing rarefaction with other normalization methods including proportions (also known as Total Sum Normalization, [TSS]), Cumulative Sum Scaling (CSS), edgeR-TMM, and DESeq-VS, TSS and rarefaction provided more accurate comparisons among communities and were the only methods that fully normalized read depths across samples (McKnight et al., 2018). In the present study, rarefying the data to 10,900 reads per sample was performed to normalize read depths across all samples according to the sample with the lowest read depth among those over 10,000 reads, while preserving sufficient read depths for each sample. Furthermore, rarefaction was chosen to meet the requirement for the source tracking analysis using both SourceTracker and FEAST algorithms, as both tools require an input ZOTU table composed of integer count values, and currently do not accept normalized ZOTU tables containing non-integer data generated by the other methods. To comprehensively compare the relative contribution of different potential sources from broad to specific category levels, spawning water, hatching water, rearing water, exchange water, and sediment were categorized as “environmental sources”. Dry feeds (including commercial spirulina powder, shrimp flakes, mixed feed, polysaccharide, and vitamin C) and fresh feeds (including algal feed, frozen *Artemia*, and live *Artemia*) were categorized as “input sources”. And environmental and input sources were considered as external sources. Bacterial communities of larvae at previous developmental stage(s) (including fertilized eggs) were also involved in source tracking analyses as internal sources (indicating internal succession). To deal with uneven sequence numbers of a given bacterial group across samples, only FEAST was used to further assess the relative contribution of above potential sources to the top 3 relatively abundant bacterial families (*Roseobacteraceae*, *Vibrionaceae*, and *Flavobacteriaceae*) of larval bacterial communities. Bacterial source tracking in the rearing water was also performed with both tools.

### 2.5 Inference of processes governing the assembly of bacteria from potential sources into larval and water communities

The number and proportion of bacterial ZOTUs being unique or shared among the major internal or external sources and shrimp larvae at a given developmental stage were shown by Venn plots using the R package ‘ggVennDiagram’ (Gao et al., 2024). The Sloan neutral model (Sloan et al., 2006) was used to infer ecological processes by which bacteria were assembled from potential sources into larval or water bacterial communities with previously reported R codes (Burns et al., 2016). At a given developmental stage, neutral model fitting was performed for larval bacterial communities considering major internal sources (including fertilized egg/larval bacterial communities from previous stage(s) that contribute more than 2% estimated by either SourceTracker or FEAST for either shrimp strain) as the species pool. And a similar model fitting was also done considering major external sources (including bacterial communities of the environmental elements and inputs that contribute more than 2%) as the species pool. The 2% threshold was applied to refine the model fitting scope, focusing on the primary sources, which collectively accounted for 94.8%-100.0% and 95.8%-100% of the known contribution based on the two source-tracking tools as described below. The goodness of model fitting was evaluated by R^2^, ranging from ≤ 0 (not fit) to 1 (perfectly fit). The 95% confidence intervals of the model were calculated by bootstrapping with 1,000 replicates. The estimated migration rate (*m*), indicating the probability of stochastic losses of individuals in the local community being replaced by dispersal from the species pool, was calculated by a nonlinear least-squares fitting using the R package ‘minpack.lm’ (Elzhov, 2016). This parameter serves as an indicator of dispersal limitation, where higher *m* values indicate lower dispersal limitation (Burns et al., 2016).

The ZOTUs within the 95% confidence intervals of the model are considered as neutrally distributed. The ZOTUs that distribute above the 95% confidence interval (above prediction) are likely positively selected by the host or have a stronger dispersal capability relative to others. The ZOTUs that distribute below the 95% confidence interval (below prediction) are selected against by the host or dispersal limited from the potential sources. The cumulative relative abundance of neutrally distributed and non-neutrally distributed (above and below prediction) ZOTUs was calculated as a metric to infer the relative influence of dispersal and drift (stochastic processes) and selection (deterministic processes) in governing the assembly of bacteria from potential sources into the community of larvae or water (Wang, Y. et al., 2020; Yan et al., 2024). Taxonomic distribution of three categories of ZOTUs at the family level was shown corresponding to each major source for each shrimp strain.

### 2.6 General statistical analysis

Alpha-diversity indices (richness of ZOTUs, Shannon-Wiener index, and phylogenetic diversity), Pielou’s evenness, and the beta-diversity metric (Bray-Curtis dissimilarity) were calculated using the R packages ‘vegan’ (Oksanen et al., 2013) and ‘picante’ (Kembel et al., 2010). Kruskal-Wallis test was applied to test the differences in bacterial alpha-diversity indices of shrimp larvae and potential bacterial sources using the R package ‘dunn.test’ (Dinno, 2014). Principal Coordinates Analysis (PCoA) was used to visualize the compositional variation of bacterial communities across shrimp larvae and potential bacterial sources using the R package ‘vegan’.

## 3 Results

### 3.1 Alpha-diversity of larval bacterial community and its potential sources

In general, bacterial alpha-diversity (including ZOTU richness, Shannon index, and phylogenetic diversity) and evenness indices of environmental sources (including spawning, hatching, rearing waters, and sediment) were either similar or higher relative to larval shrimps across developmental stages, while input sources showed similar (dry feed and algal feed) or lower (especially frozen/live *Artemia*) bacterial alpha-diversity compared with the host (Supplementary Fig. 1).

### 3.2 Bacterial community compositions of the elements in the larviculture system

Principal coordinates analysis (PCoA) showed that bacterial community compositions (based on ZOTUs) were distinctly clustered according to larval shrimps and different categories of elements in the larviculture system as potential bacterial sources, despite large within-group variance for larval shrimps and rearing waters (Fig. 2). Although bacterial communities of fertilized egg and larval shrimps were overall predominated by *Alphaproteobacteria* (primarily the families *Roseobacteraceae* and *Paracoccaceae*), *Gammaproteobacteria* (primarily the families *Vibrionaceae*, *Alteromonadaceae*, *Pseudoalteromonadaceae*, and *Halothiobacillaceae*), and *Bacteroidota* (primarily the families *Flavobacteriaceae* and *Weeksellaceae*), regardless of SR and FG strains (accounting for 94.2% and 94.1% of relative abundance in average, respectively), clear succession of taxonomic composition of larval bacterial community was observed across host developmental stages, especially at the family level (Supplementary Fig. 2).

**Figure 2.**
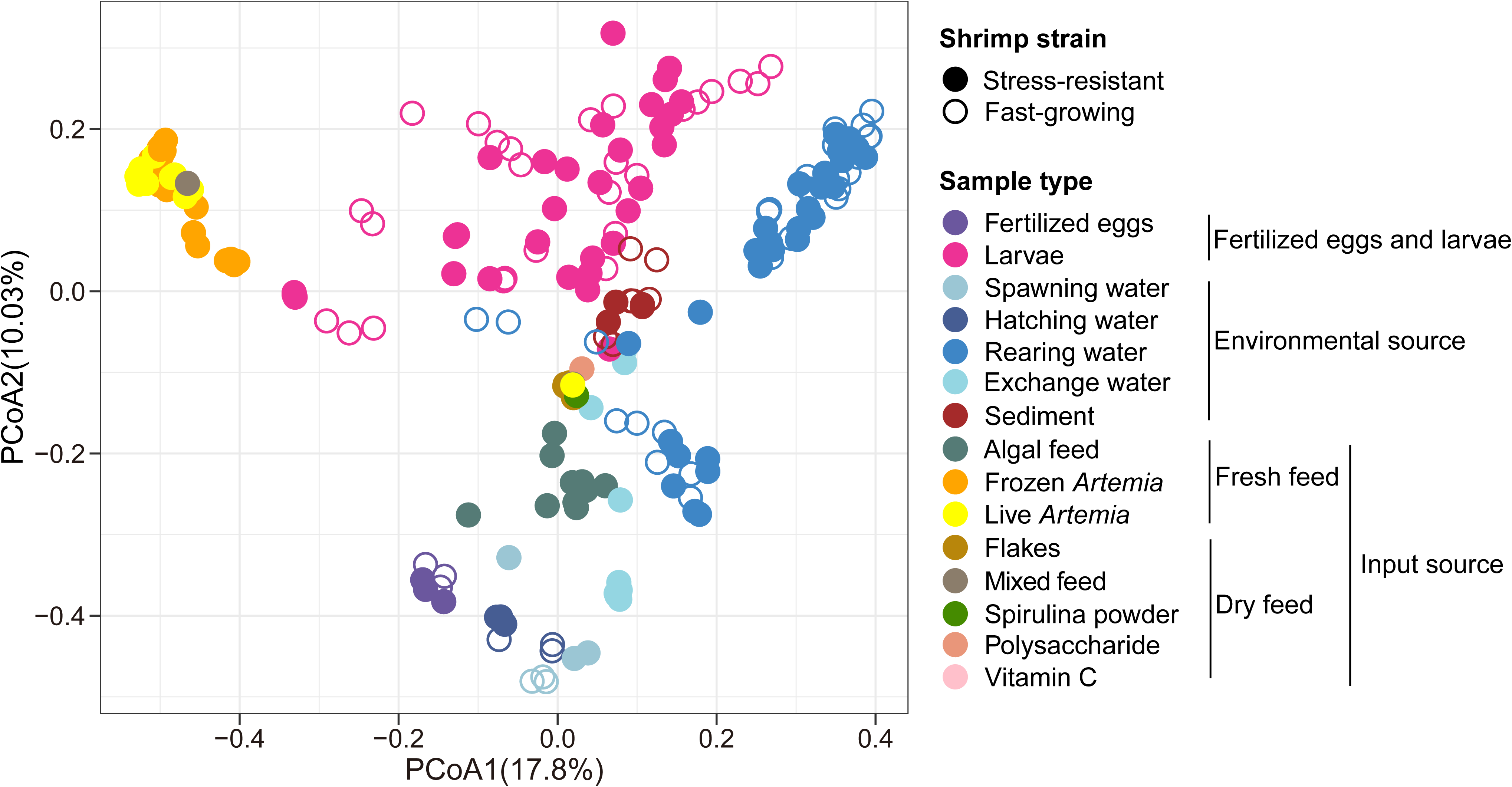
Principal Coordinate Analysis (PCoA) based on Bray-Curtis dissimilarity visualizing the compositional variation of bacterial communities across fertilized eggs, shrimp larvae, and environmental and input sources during the larviculture.

For environmental elements, bacterial communities in the spawning, hatching, rearing, exchange waters were overall dominated by *Alphaproteobacteria*, *Gammaproteobacteria*, and *Bacteroidota*, regardless of ponds culturing shrimp strain (Supplementary Fig. 3). *Actinomycetota* became abundant in the rearing waters from *zoea* II to *postlarvae* V. At the family level, bacterial taxonomic composition in the rearing water at the beginning of larviculture (*naupilus* I) was similar to the exchange waters added at the later stages (*mysis* I and *postlarvae* I), with dominance of *Oceanospirillaceae*, *Roseobacteraceae*, *Vibrionaceae, Haliscomenobacteraceae*, *and Pelagibacteraceae* quickly switched to be overwhelmingly dominated by *Roseobacteraceae* after larval mouth opening at *zoea* I (reaching up to 14.1%-59.1% during *zoea* II to *postlarvea* V). The *Bacteroidota* families *Haliscomenobacteriaceae* and *Flavobacteriaceae* became increasingly dominant in the bacterial community of rearing waters during the mouth-opening stages of larvae (*nauplius* VI - *zoea* I and *zoea* I) and remained at considerable levels at the later stages. In addition, *Vibrionaceae* was commonly abundant in the rearing waters during the mouth-opening stages of larvae.

Distinct combinations of feeds were input into the larviculture system to meet the needs of larval development and growth at different stages. We found that the taxonomic composition of bacteria largely varied across different types of inputs (Supplementary Fig. 4). For fresh feeds, bacterial communities of algal feed were dominated by *Alphaproteobacteria* (28.1% in average; mainly *Roseobacteraceae* (16.6%)) and *Bacteroidota* (52.6%; mainly *Flavobacteriaceae* (41.5%)), while it is worth noting that bacterial communities of *Artemia* (frozen and live) were overwhelmingly dominated by *Gammaproteobacteria* (mainly *Vibrionaceae*, accounting for 69.0% and 57.3% in frozen and live *Artemia*). Bacterial communities of dry feeds vary with kind, with spirulina powder dominated by *Cyanobacteriota* (89.6%, mainly *Sirenicapillariaceae*), shrimp flakes (1^#^-5^#^) dominated by *Bacillota* (60.1%, mainly *Bacillaceae* and *Lactobacillaceae*) and *Cyanobacteriota* (29.4%, mainly *Sirenicapillariaceae*), mixed feed (a) dominated by *Gammaproteobacteria* (74.6%, mainly *Vibrionaceae* and *Alteromonadaceae*), and mixed feed (b) dominated by *Lactobacillaceae* (91.5%).

### 3.3 Relative contribution of potential sources to the bacterial community and major taxonomic groups of larval shrimps

SourceTracker and FEAST generally showed similar trends in tracking larval bacterial sources at most developmental stages, regardless of shrimp strains (Fig. 3). Specifically, bacteria of naupliar shrimps (*nauplius* I) before introduced into the rearing water was considerably from hatching water and fertilized eggs (accounting for 24% and 41.2% of contribution as estimated by SourceTracker and FEAST); however, the majority of sources remain unknown (74% and 58.7% by SourceTracker and FEAST). During the transitional stage from *nauplius* VI to *zoea* I, SourceTracker revealed that rearing water was the major source of larval bacterial communities (73%-89%), and larvae from the previous stage (*nauplius* I) also contributed to some extent; although FEAST confirmed the importance of these two sources, large proportion of sources remains unknown. After larval mouth opening at *zoea* I, larvae from the previous stage(s) (indicating internal succession) became the most important larval bacterial source regardless of source-tracking tools or shrimp strains, and the predominance of internal succession was overall strengthened with host development (from *zoea* sub-stages (63.2%-63.5%) to *mysis* sub-stages (75.9%-81.8%) then to *postlarvae* sub-stages (93.9%-94.2%)). Despite this main trend, environmental (rearing water) and input (mainly frozen *Artemia* and/or dry feed) sources also considerably contributed to the larval bacterial community at *zoea* II-III and *mysis* I. Overall, no major shrimp-strain-specific features were detected, expect differences in relative contribution of sources at sub-category level, as shown in the inner ring of Fig. 3 (eg. among larvae from different previous stages). Subtly, algal feed only contributed a bit during the mouth-opening stages (*nauplius* VI - *zoea* I and *zoea* I) of SR strain.

**Figure 3.**
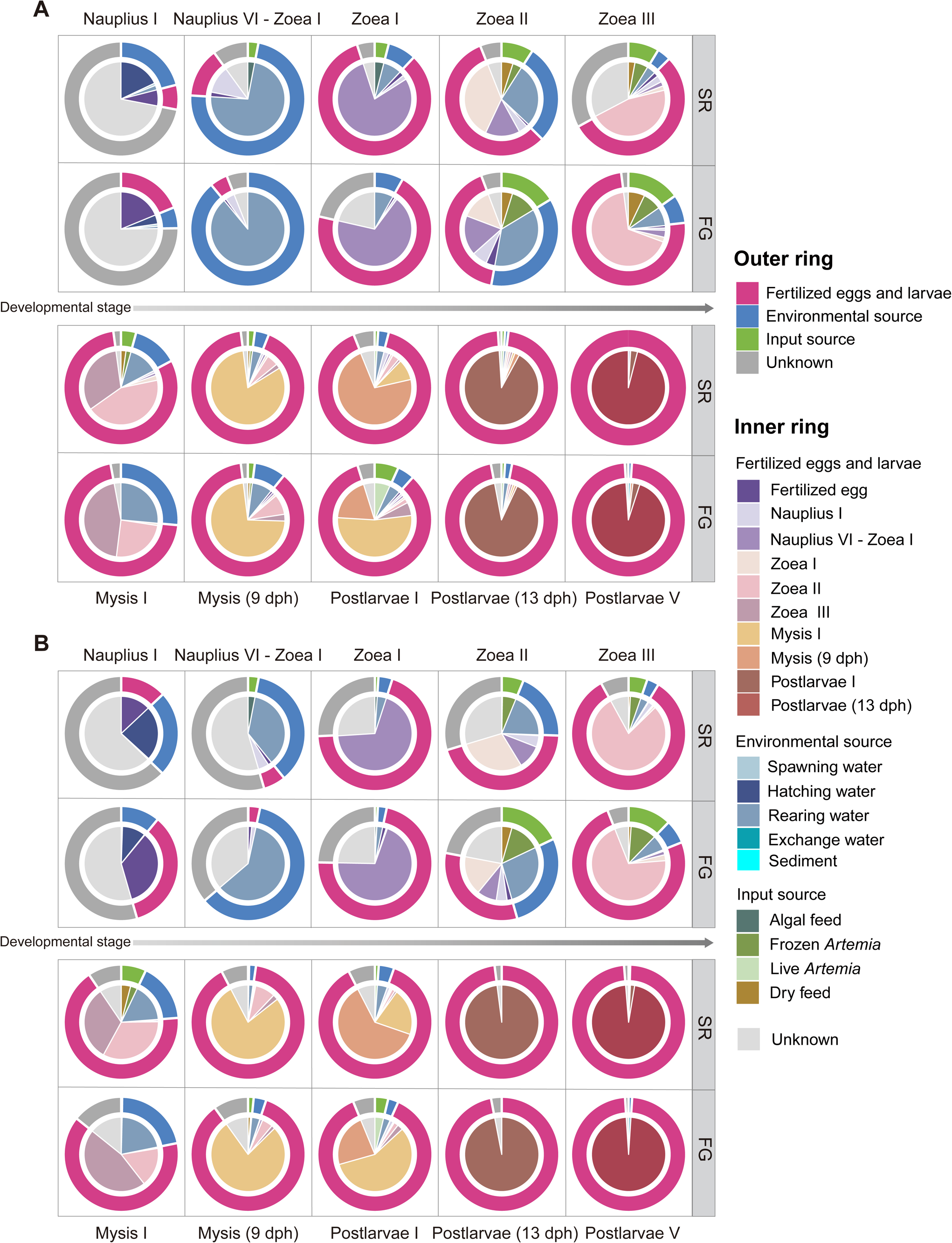
The relative contribution of various potential sources to the bacterial community of shrimp larvae across host developmental stages inferred by SourceTracker (A) and Fast Expectation-maximization for Microbial Source Tracking (FEAST; B). SR, stress-resistant shrimp strain; FG, fast-growing shrimp strain.

Despite the dynamics of taxonomic composition of larval bacterial community with host development, as the top 3 relatively abundant bacterial families, *Roseobacteraceae* was overall predominant across developmental stages; *Vibrionaceae* was more abundant during *nauplius* I to *zoea* III (especially at *nauplius* VI - *zoea* I and *zoea* I) than for the later stages, while *Flavobacteriaceae* tended to be more abundant at *mysis* and *postlarvae* than for the former stages (Supplementary Fig. 2). Regardless of shrimp strains, *Roseobacteraceae* assemblages of naupliar shrimps (*nauplius* I) were mainly from fertilized eggs and environmental source (including hatching, spawning, and rearing waters) with nearly equal contributions, while rearing water became a major source of *Roseobacteraceae* assemblages at the transitional stage from *nauplius* VI to *zoea* I, with a considerable faction remains unknown (Fig. 4A). Similar to the whole bacterial community, internal succession from previous stage(s) became the most important *Roseobacteraceae* source of larval shrimps (except *zoea* II, when rearing water contributed more than internal source), along with the predominance of internal source strengthened with host development. As a bacterial group with many opportunistic pathogenic members, *Vibrionaceae* assemblages of larval shrimps were mainly contributed by the rearing water and/or internal source during *nauplius* I to *zoea* I, and then frozen/live *Artemia* became the second most important following internal source during *zoea* II to *postlarvae* (13 dph), when environmental sources barely contributed, except that the exchange water made a contribution to SR strain at *postlarvae* I (Fig. 4B). Hatching water was the predominant source of *Flavobacteriaceae* assemblages of naupliar shrimps (*nauplius* I), and internal succession became the most important source of *Flavobacteriaceae* assemblages during *nauplius* VI - *zoea* I to *postlarvae* V (Fig. 4C). In addition, environmental (rearing water) and/or input (frozen/live *Artemia*) sources commonly contributed to larval *Flavobacteriaceae* assemblages and often showed between-shrimp-strain differences.

**Figure 4.**
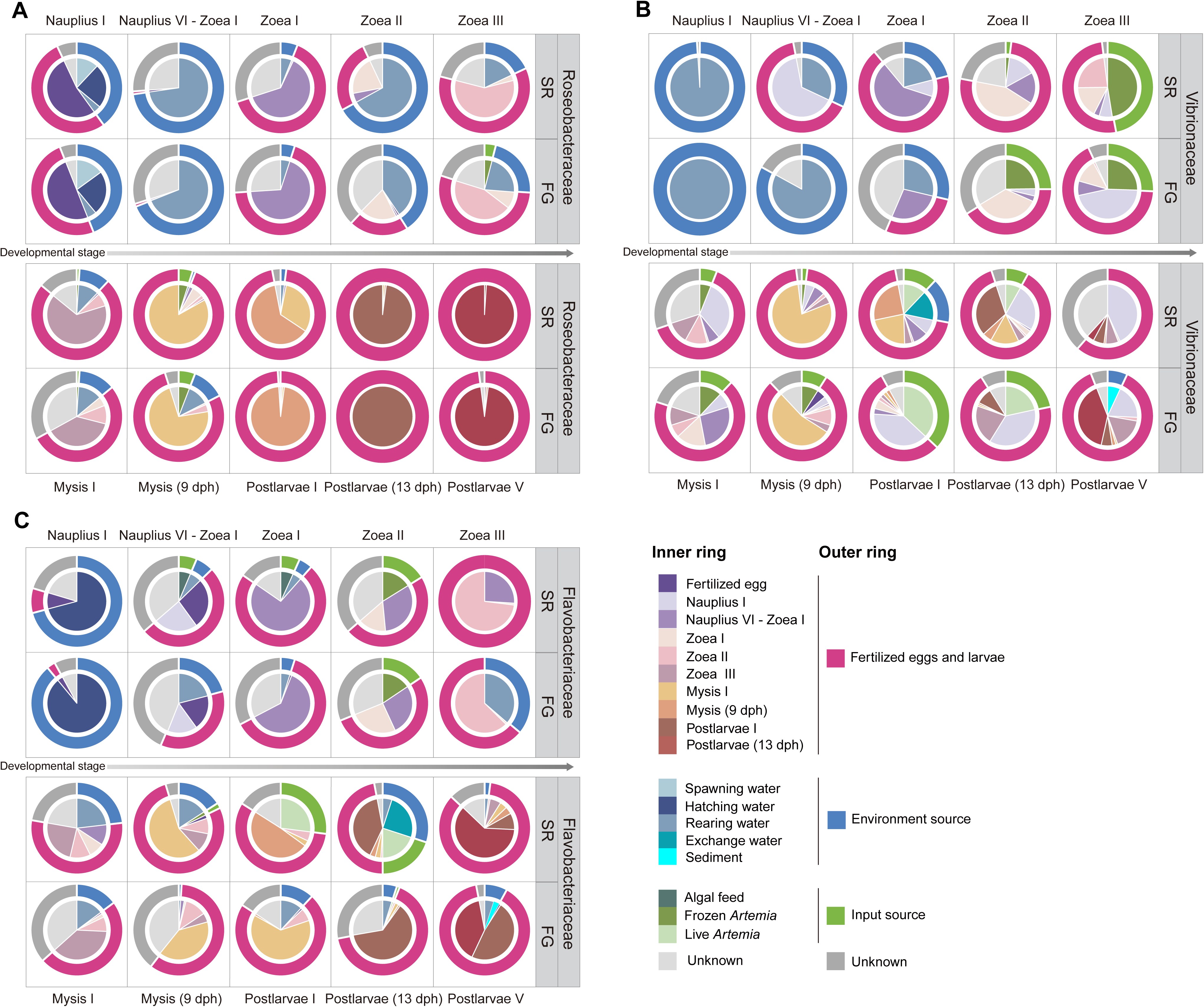
The relative contribution of various potential sources to top 3 abundant bacterial families (*Roseobacteraceae*, *Vibrionaceae*, and *Flavobacteriaceae*) of shrimp larvae across host developmental stages inferred by Fast Expectation-maximization for Microbial Source Tracking (FEAST). To ensure the reliability of source tracking, the samples of larvae and potential sources with sequences greater than 1,000 for each family were retained for analysis, and thus the rearing water samples and the spawning water samples were excluded from the source tracking of *Vibrionaceae* (from *zoea* II to *postlarvae* V) and *Flavobacteriaceae* (*nauplius* I), respectively. SR, stress-resistant shrimp strain; FG, fast-growing shrimp strain.

### 3.4 Relative contribution of potential sources to the bacterial community in rearing waters

Like larval bacterial community, both tools showed overall similar performances on tracking bacterial sources of rearing waters at most developmental stages (especially after *zoea* II) for both shrimp strains (Supplementary Fig. 5). At *nauplius* I, the majority of sources of bacteria in the rearing water were unknown, though naupliar shrimps contributed a bit, as estimated by SourceTracker, while naupliar shrimps contributed more and a large proportion of sources remains unknown as estimated by FEAST. During the mouth-opening stages of larvae (*nauplius* VI - *zoea* I and *zoea* I) and afterwards, rearing waters from previous stage(s) became the most important source of bacterioplankton regardless of shrimp strains or tools (from *zoea* sub-stages (26.0%-96.1%) to *mysis* sub-stages (91.4%-93.3%) then to *postlarvae* sub-stages (90.8%-96.6%)), suggesting that succession is the main bacterial source of rearing water. During *nauplius* VI - *zoea* I to *zoea* II, larval shrimps commonly made certain contribution to bacterioplankton in rearing waters, despite the differences between shrimp strains or tools. Input source (algal feeds) only contributed to bacterioplankton at *nauplius* VI - *zoea* I and *zoea* I. Other fresh feeds and all dry feeds barely contributed to bacterioplankton at any stages.

### 3.5 Ecological processes governing the assembly of bacteria from major sources into the shrimp larval community

The proportion of bacterial ZOTUs that larvae shared with all major internal sources ranged from 41.41% to 71.24% across host developmental stages (Fig. 5). When assuming the major internal sources as the species pool, the occurrence of the ZOTUs shared with the source(s) and the host either did not (at the beginning (*nauplius* I) and ending (*postlarvae* (13 dph and V) stages for both strain and *postlarvae* I for FG strain as well) or barely/weakly fit the neutral models (Fig. 5). Regardless of shrimp strains or developmental stages, the above-prediction ZOTUs overwhelmingly dominated the accumulative relative abundance of ZOTUs shared with all or each internal source(s) (*nauplius* I: 98.78%-98.83%, *nauplius* VI - *zoea* I: 96.55%-98.40%, *zoea*: 93.13%-98.02%, *mysis*: 94.22%-98.39%, and *postlarvae*: 94.67.0%-98.74%), suggesting most proportion of bacteria potentially from internal source(s) were deterministically succeeded by the host.

**Figure 5.**
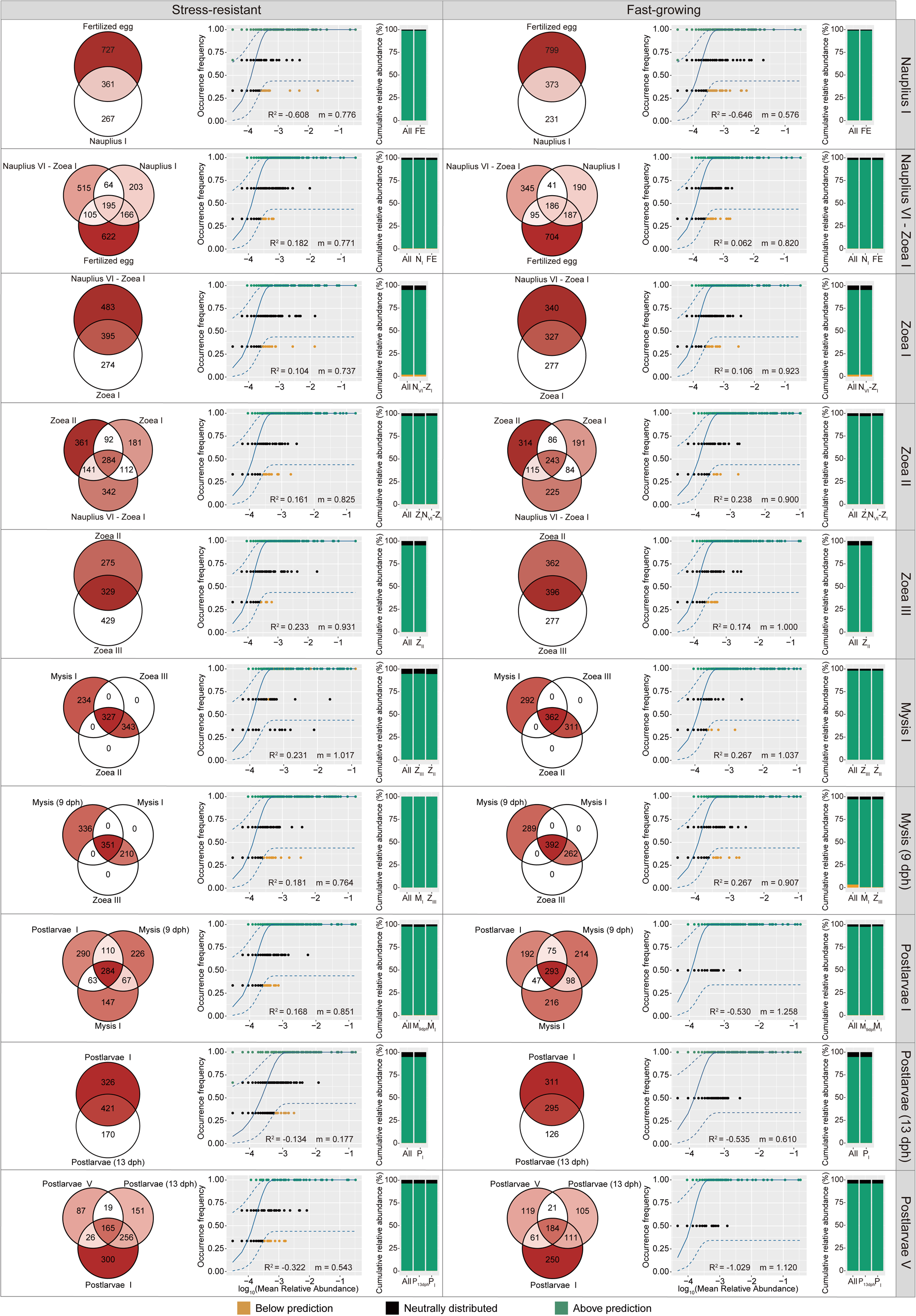
Venn diagrams show the number and proportion (as indicated by the shade of the color; darker colors correspond higher proportions) of bacterial ZOTUs being unique or shared among the major internal sources (larval bacterial communities from previous stages that contribute more than 2% at a given developmental stage estimated by either SourceTracker or FEAST, regardless of shrimp strains) and shrimp larvae at a given developmental stage. Fitting of the neutral models for shrimp larval bacterial communities with the major internal sources (the shared fraction) as the species pool. The goodness of model fitting was evaluated by R^2^, ranging from ≤ 0 (not fit) to 1 (perfectly fit). The bacterial ZOTUs that occur more frequently than the model prediction is shown in green, those occur less frequently than predication is shown in orange, those fit the neutral distribution is shown in black. Blue dashed lines represent 95% confidence intervals around the model prediction. The histograms show the cumulative relative abundance of three categories of ZOTUs (above prediction, below prediction, and neutrally distributed) from all major internal sources (All) or each source (larvae from previous stage(s)). N_I_, *nauplius* I; N_VI_-Z_I_, *nauplius* VI - *zoea* I; Z_I_, *zoea* I; Z_II_, *zoea* II, Z_III_, *zoea* III; M_I_, *mysis* I; M_9dph_, *mysis* (9 dph); P_I_, *postlarvae* I; P13dph, *postlarvae* (13 dph); P_V_, *postlarvae* V.

The proportion of ZOTUs that larvae shared with all major external sources ranged from 3.64% to 53.03% (Fig. 6). When assuming the major external sources as the species pool, the occurrence of the ZOTUs shared with the source(s) and the host did not fit the neutral models for both shrimp strains at all stages (Fig. 6). Similar to internal sources, the above-prediction ZOTUs predominated the accumulative relative abundance of ZOTUs shared with all or each major external source(s) (*nauplius* I: 98.86%-99.40%, *nauplius* VI - *zoea* I: 70.03%-94.70%, *zoea*: 60.00%-99.49%, *mysis*: 64.86%-98.33%, and *postlarvae* I: 76.81%-99.76%), suggesting that larval shrimps exhibited a strong preference for deterministic selection towards external bacteria.

**Figure 6.**
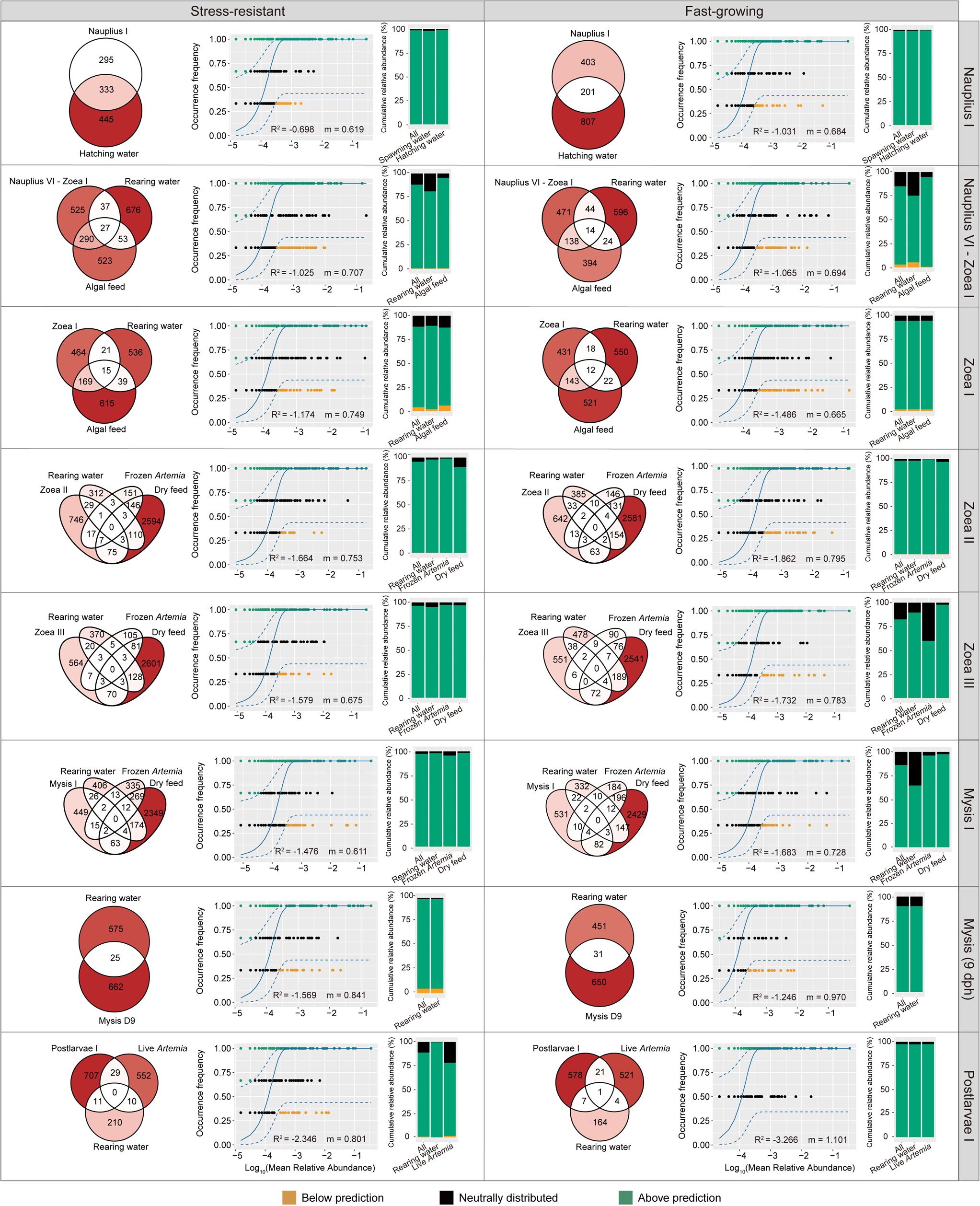
Venn diagrams show the number and proportion (as indicated by the shade of the color; darker colors correspond higher proportions) of bacterial ZOTUs being unique or shared among the major external sources (bacterial communities from environment and inputs that contribute more than 2% at a given developmental stage estimated by either SourceTracker or FEAST, regardless of shrimp strains.) and shrimp larvae at a given developmental stage. Fitting of the neutral models for shrimp larval bacterial communities with the major external sources (the shared fraction) as the species pool. At the stages *postlarvae* (13 dph) and *postlarvae* V, no external source contributed more than 2%, and thus no modeling was performed for these two stages. The goodness of model fitting was evaluated by R^2^, ranging from ≤ 0 (not fit) to 1 (perfectly fit). The bacterial ZOTUs that occur more frequently than the model prediction is shown in green, those occur less frequently than predication is shown in orange, those fit the neutral distribution is shown in black. Blue dashed lines represent 95% confidence intervals around the model prediction.

### 3.6 Taxonomic selection of bacteria by larval shrimps from potential sources

To reveal which bacterial taxa shrimps preferred to succeed or selected from major internal or external sources, taxonomic distribution of three categories of ZOTUs assessed by the neutral model was shown corresponding to internal or external sources for each shrimp strain (Figs. 7-8). According to taxonomic distribution of ZOTUs above prediction, the two shrimp strains showed overall similar taxonomic preferences when succeeding bacteria internally from previous stage(s), during *nauplius* I to *zoea* II and *postlarvae* sub-stages, though subtle between-shrimp-strain differences could be found at stages like *nauplius* VI - *zoea* I, *zoea* III, and *mysis* sub-stages (Fig. 7). At *nauplius* I - *zoea* I, both strains preferred to succeed *Vibrionaceae*, *Roseobacteraceae*, and *Paracoccaceae* taxa, while SR strain also succeeded *Halomonadaceae*, *Flavobacteriaceae*, and *Haliscomenobacteraceae* taxa. At *zoea* III, in addition to the preference of *Weeksellaceae*, *Vibrionaceae*, *Roseobacteraceae*, *Colwelliaceae*, and *Flavobacteriaceae* taxa by both strains, FG strain also showed a preference for *Vicingaceae*, *Pseudomonadaceae*, and *Paracoccaceae* taxa. At *mysis* I, both strains preferred to succeed *Roseobacteraceae*, *Flavobacteriaceae*, *Paracoccaceae*, *Vibrionaceae*, and *Weeksellaceae* taxa, while FG strain uniquely took *Vicingaceae*, *Halieaceae*, and *Pirellulaceae* taxa into preferences. At *mysis* (9 dph), similar preferences for *Roseobacteraceae*, *Flavobacteriaceae*, *Paracoccaceae*, and *Cohaesibacteraceae*, and *weeksellaceae* taxa by both strains were shown, as well as specific preferences for *Marinobacteraceae*, *Pseudoalteromonadaceae*, and *Halothiobacillaceae* taxa by SR strain, and for *Devosiaceae*, *Moraxellaceae*, and *Vicingaceae* taxa by FG strain.

**Figure 7.**
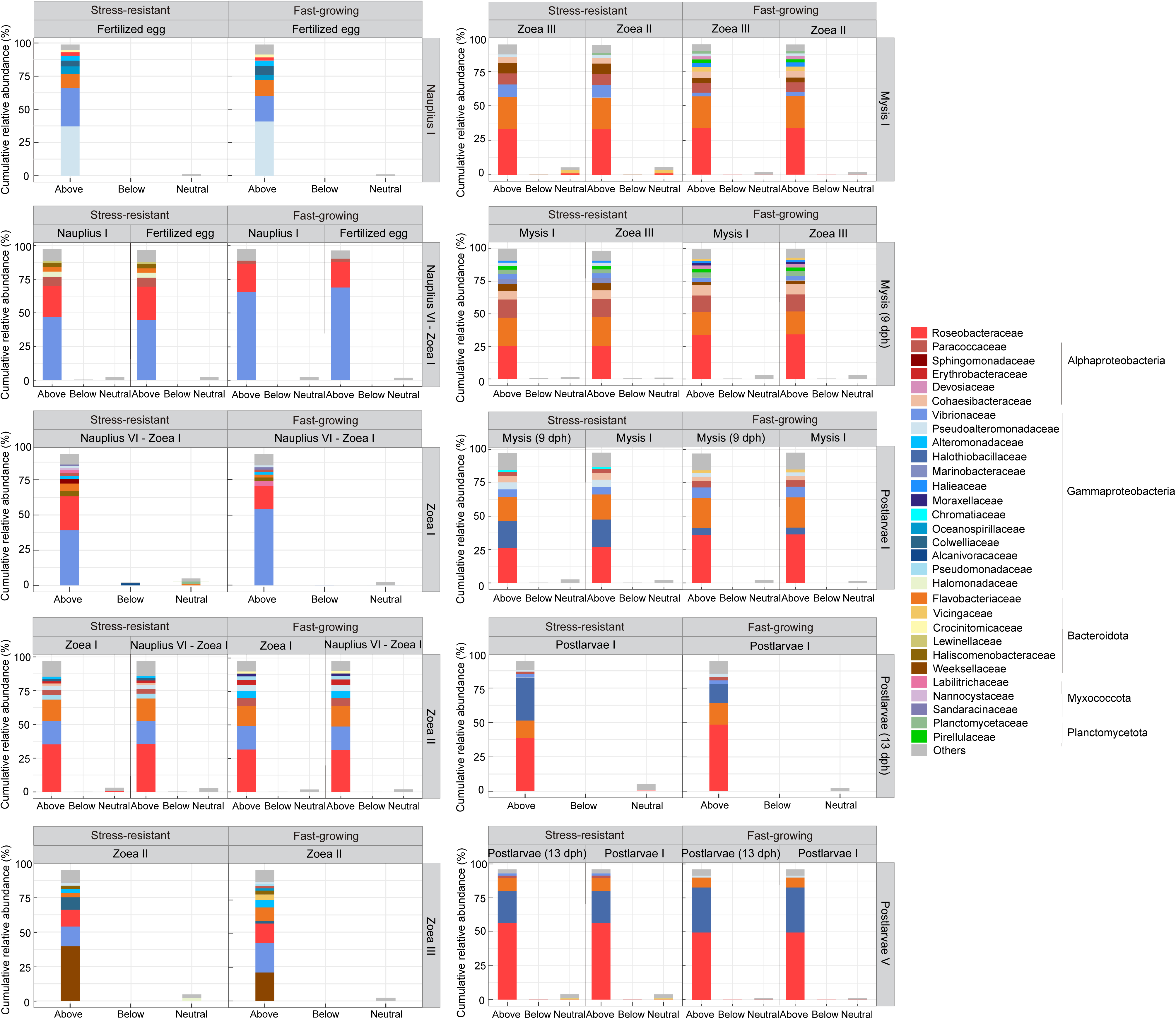
The cumulative relative abundance and taxonomic distribution of three categories of shared ZOTUs (relative abundances >1%) (above prediction (Above), below prediction (Below), and neutrally distributed (Neutral)) between each internal source and larvae at a given developmental stage inferred by the neutral models in Fig. 5

**Figure 8.**
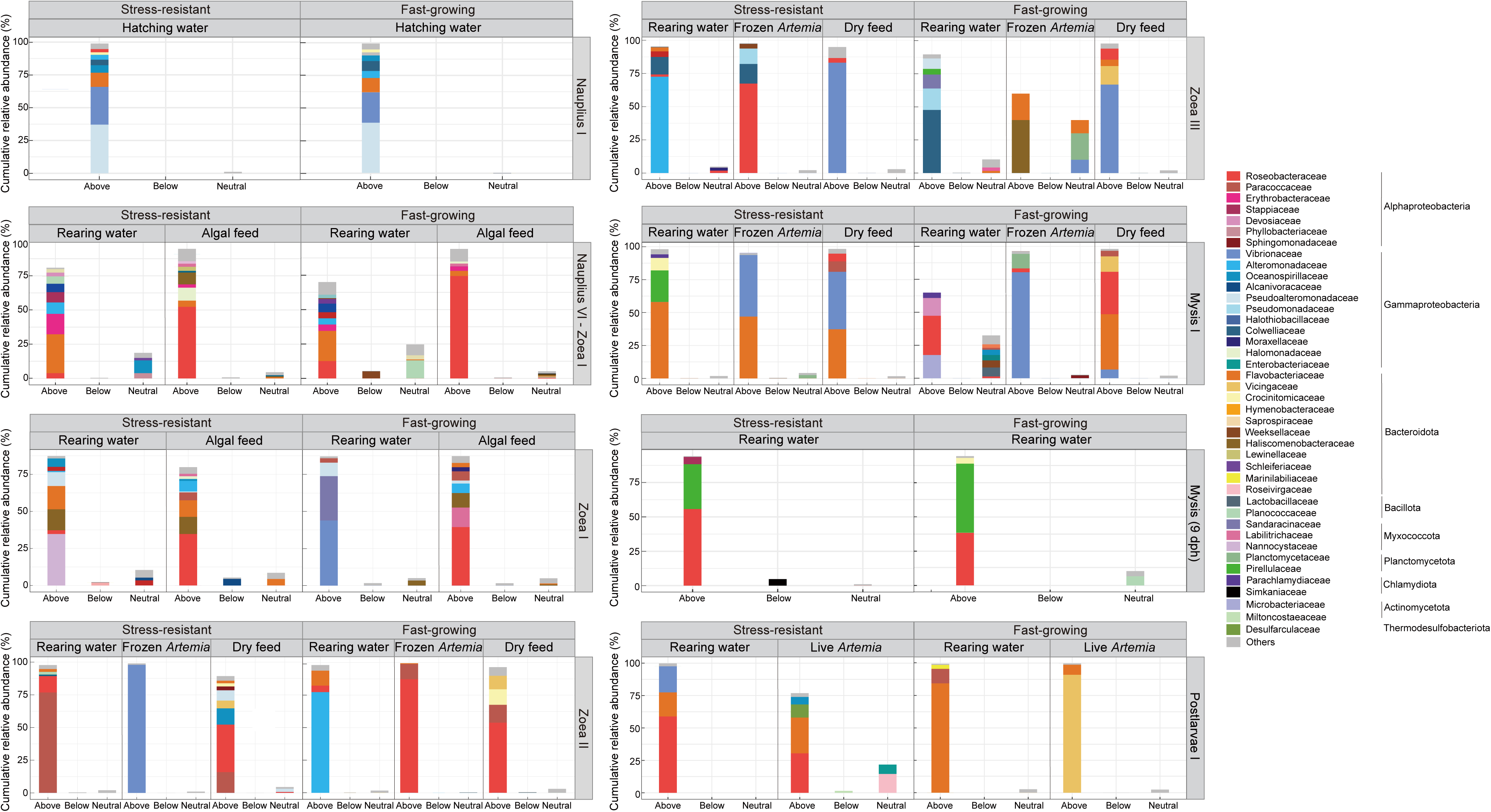
The cumulative relative abundance and taxonomic distribution of three categories of shared ZOTUs (relative abundances >1%) (above prediction (Above), below prediction (Below), and neutrally distributed (Neutral)) between each external source and larvae at a given developmental stage inferred by the neutral models in Fig. 6.

Unlike the internal succession, the two shrimp strains commonly exhibited respective preferences in selecting for external bacteria (when they mattered) since the mouth of larvae began to open (Fig. 8). Specifically, at the transitional stage of *nauplius* VI - *zoea* I, SR strain tented to select higher proportion of bacteria from the rearing water than for FG strain but with similar taxonomic composition (mainly *Flavobacteriaceae*, *Erythrobacteraceae*, *Alteromonadaceae*, *Stappiaceae*, and *Desulfuromonadaceae*), and both strains (especially FG) preferred to select *Roseobacteraceae* taxa from algal feed, from which SR strain also selected *Haliscomenobacteraceae* and *Halomonadaceae* taxa. At *zoea* I, SR strain preferred to select *Nannocystaceae*, *Flavobacteriaceae*, *Haliscomenobacteraceae*, *Oceanospirillaceae*, and *Roseobacteraceae* taxa from water, from which FG strain preferred to select *Vibrionaceae*, *Sandaracinaceae*, and *Pseudoalteromonadaceae* taxa; both strains preferred to select *Roseobacteraceae*, *Haliscomenobacteraceae*, and *Alteromonadaceae* taxa from algal feed. At *zoea* II, SR strain preferred to select *Paracoccaceae* and *Roseobacteraceae* taxa from water and *Vibrionaceae* taxa from frozen *Artemia*, while FG strain primarily selected *Alteromonadaceae* and *Flavobacteriaceae* taxa from water and *Roseobacteraceae* and *Paracoccaceae* taxa from frozen *Artemia.* At *zoea* III, SR strain preferred to select *Alteromonadaceae* and *Colwelliaceae* taxa from water, from which FG strain specifically selected *Colwelliaceae*, *Pseudomonadaceae*, and *Sandaracinaceae* taxa; meanwhile, SR strain selected *Roseobacteraceae*, *Pseudomonadaceae*, and *Colwelliaceae* taxa but FG strain selected *Haliscomenobacteraceae* and *Flavobacteriaceae* taxa from frozen *Artemia*, from which *Planctomycetaceae*, *Flavobacteriaceae*, and *Vibrionaceae* taxa also entered FG strain *via* neutral processes. At *mysis* I, the two strains selected largely different taxa from water (*Flavobacteriaceae*, *Pirellulaceae*, and *Crocinitomicaceae* for SR strain and *Roseobacteraceae*, *Devosiaceae*, and *Microbacteriaceae* for FG strain); SR strain selected *Vibrionaceae* and *Flavobacteriaceae* taxa from frozen *Artemia*, from which FG strain selected *Vibrionaceae* and *Planctomycetaceae* taxa. At *mysis* (9 dph), both strains similarly selected *Pirellulaceae* and *Roseobacteraceae* taxa from water. At *postlarvae* I, SR strain selected *Roseobacteraceae*, *Vibrionaceae*, *Flavobacteriaceae* taxa from water and *Roseobacteraceae*, *Flavobacteriaceae*, and *Desulfarculaceae* taxa from live *Artemia*, while FG strain selected *Flavobacteriaceae* and *Paracoccaceae* taxa from water and *Vicingaceae* taxa from the live *Artemia*.

## 4 Discussion

### 4.1 The mouth opening is a turning point in bacterial source composition of larvae

Regardless of shrimp strains or source-tracking tools, source compositions of larval bacterial communities can be generally divided into two major phases: before and after the mouth opening of larval shrimps, going through a transition from external-source-dominated to internal-succession-dominated. Before the mouth opening, it is reasonable that naupliar bacterial community (*nauplius* I) was derived from the vertical transfer from fertilized eggs and the attachment of bacterioplankton in the hatching ponds. However, the major proportion of bacterial sources of nauplii (*nauplius* I) before being introduced into the rearing water remains unknown. The unknown sources may include the carrying water (which was not sampled) during the transfer of naupliar shrimps after the hatching (for ∼ 1.5 h), indicating that the carrying water could be a potential but neglected juncture in establishing naupliar microbiota. The large contribution of the rearing water to the larval bacterial community at *nauplius*-VI-to-*zoea*-I transitional stage and the strong internal succession afterwards for both larval and water bacterial communities suggest that the bacterial community in rearing water at the beginning of larviculture likely has a persistent impact on larval bacterial community later, which should be regulated into a beneficial status by initial treatment of rearing water before introducing nauplii. These observations provide a theoretical basis for the empirical importance of “the cultivation of water” in larviculture practice (Gundersen et al., 2022; Hansen and Olafsen, 1999), though “what is a beneficial status of rearing water” remains an open question. When tracing bacterial sources in the rearing water, it was found that the main sources of bacterioplankton before larval mouth opening remain unknown, speculatively from the original coastal water. This further emphasizes the importance of initial cultivation of rearing water in initiating and maintaining a healthy microbiota in the rearing water, that is, targeted reestablishment of microbiota after filtration and disinfection of the original coastal water. After the mouth opening, there was only a very short period (*nauplius* VI - *zoea* I and *zoea* I) when algal feed made a contribution to bacterioplankton in the rearing water (regardless of tools or shrimp strains), suggesting that feed input exhibited limited impact on bacterioplankton community, likely due to their quick consumption by predation of larvae. On the other hand, a previous study based on the absolute abundance of bacteria demonstrated that 34% of the rearing water bacterial community over the entire larviculture were introduced by algal feed, considering the growth of the introduced taxa in the rearing water (Heyse et al., 2021). This accumulative effect on early introduced taxa to later stages might be considered as succession in our present study. Bacterioplankton communities in the rearing waters at *zoea* sub-stages (especially *zoea* II) showed a signature of receiving a certain contribution from larval shrimps and vice versa as discussed below, suggesting frequent water-larva exchange of bacteria after larval mouth opening. Meanwhile, considering the less or negligible contribution from feed input, the best practice for microbial regulation during larviculture should be conducted through the rearing water (as the best pathway) as approaching *zoea* and at *zoea* (especially *zoea* II, as the best timing).

After the mouth opening, internal succession became the main source of the larval bacterial community (except *zoea* II), confirming the criticality and potential persistence of initial establishment of microbiota in the early life of aquatic invertebrates, as similar to that observed in mammals (Bi et al., 2019; Kerr et al., 2015). This pattern could be explained by priority effect/colonization resistance driven by niche occupation of bacterial commensals in the digestive tract of larvae after the mouth opening, which built barriers against external invaders (De Schryver and Vadstein, 2014; Xiong et al., 2019). It is worth noting that external sources still made non-negligible contribution to the larval bacterial community during *zoea* sub-stages, especially at *zoea* II, when external sources (mainly rearing water, frozen *Artemia*, and/or dry feeds, also including unknown sources logically presumed to be external) made similar or even greater contribution than internal succession. The trend that larval bacterial communities at three *zoea* sub-stages were more influenced by bacterioplankton in the rearing water relative to the later stages has been indicated by our previous work (Wang, Y. et al., 2020), and here we further quantitatively confirmed this pattern *via* source tracking. This suggests that zoeal shrimps (especially at *zoea* II) are somehow vulnerable to the invasion of external bacteria, likely due to the shift of nutrient acquisition mode to predation and the immature digestive tract, making it easier for external bacteria in rearing water or feeds to migrate and then to colonize (Angthong et al., 2023; Pérez-Morales et al., 2017). The dominance of external sources at *zoea* II suggests the importance of ensuring microbial safety after larvae open their mouths and the challenge of remaining a beneficial microbial status and reducing pathogenic risk in the environment and inputs at this vital sub-stage, when larvae are physiologically vulnerable and commonly suffered from *zoea*-II syndrome, characterized by reduction in feeding rate and impairment in metamorphosis followed by high mortalities (Kumar et al., 2017). The susceptibility of larval shrimps to the invasion of external bacteria may also related to *zoea*-II syndrome and the causality between them is worthy of follow-up studies. On the other hand, the mouth-opening stage and subsequent days are likely the best opportunity to introduce proper regulation of beneficial bacteria from external sources to establish dominance in larvae. From 9 dph (*mysis*) onward, external sources showed less or negligible contribution to larval bacterial communities. Although the constant feeding led to frequent external bacterial input, the input sources only made considerable contribution to larval bacterial community for both shrimp strains at *zoea* II and III (also contributed to SR community at *nauplius* VI - *zoea* I, *zoea* I, and *mysis* I; to FG community at *postlarvae* I; and to both communities at mysis (9 dph)), suggesting that microbial regulation *via* input is highly stage-dependent. Compared with fresh feeds, most microorganisms from dry feeds are presumed to be non-viable or at least less active, thus contributing overall less to the larval bacterial community.

### 4.2 Source tracking provides guidelines for regulating potential benefit and pathogenic bacterial groups

Besides tracking larval bacterial sources at the community level, estimating major sources of key bacterial groups that play potential roles in digestive tract functioning, health maintenance, and pathogenic risk can provide a more specific and finer perspective on oriented management for beneficial host microbiota and pathogen prevention. In the currently limited studies on microbiome of *P. vannamei* larvae, despite differences in nucleic acid extraction methods, *Alphaproteobacteria*, *Gammaproteobacteria*, and *Bacteroidota* were found to be the main dominant groups of the larval bacterial community, and the *Alphaproteobacteria* family *Rhodobacteraceae* appears to be the most dominant bacterial group (Dong et al., 2023; Rungrassamee et al., 2014), which is corresponding to our founding that *Roseobacteraceae* (formerly *Rosebacter* group of the family *Rhodobacteraceae*, also the majority of *Rhodobacteraceae* in shrimp), *Vibrionaceae* (*Gammaproteobacteria*), and *Flavobacteriaceae* (*Bacteroidota*) as the top 3 most abundant families of larval bacterial community (accounting for 56.2% of sequences in the dataset). The higher relative abundance of *Rhodobacteraceae* in the gut bacterial community of shrimp has been found associated with normal growth (Dong et al., 2023; Guo et al., 2022), pathogen antagonism (Rajeev et al., 2024), cold-stress resistance (Liu et al., 2019), and thus better culture performance. Our previous work has demonstrated that glucose addition (as a basic regulatory measure in the biofloc-based culture system) improved the culture performance of shrimps by regulating the assembly of *Rhodobacteraceae* taxa in gut bacterial community. Recent studies also paid attention to the members of *Roseobacteraceae* (such as *Ruegeria* sp. and *Phaeobacter* sp.) for their potential to serve as probiotics in aquaculture due to their ability to produce tropodithietic acid, which inhibits pathogenic vibrios (Sonnenschein et al., 2017). Furthermore, most members of *Rhodobacteraceae* can synthesize vitamin B_12_ (Sanudo-Wilhelmy et al., 2014), which is essential for shrimps (Kim et al., 2022). Therefore, source tracking of *Roseobacteraceae* taxa in larval bacterial community can facilitate regulatory and isolating strategies for *Roseobacteraceae* taxa those are beneficial for larval health. It seems that the rearing water largely reshaped the initial *Roseobacteraceae* assemblages of larvae at the *nauplius*-VI-to-*zoea*-I transitional stage and kept making considerable contribution besides internal succession until *mysis* I, suggesting that microbial regulation like introducing probiotics derived from *Roseobacteraceae* should be performed as early as pre-mouth-opening stage and continue until *zoea* III or *mysis* I through the rearing water. Many practices of probiotic amendment in aquaculture were conducted by mixing probiotics into the feeds and intended to facilitate the capture of probiotics by the host with feeding (Bernal et al., 2016). However, the minor contribution from the feeds to either larval or water bacterial community at most stages, as we discussed above, suggests that directed input to the rearing water could be a more efficient way to introduce probiotics to the larviculture system. It is worth noting that a great proportion of *Roseobacteraceae* source at the early stages remains a mystery, indicating that the high diversity of species, sources, and ecological roles of this group in the larviculture system needs to be unveiled by further efforts.

As another bacterial family that reported to be positively associated with shrimp health (Huang et al., 2020), *Flavobacteriaceae* hold potentials in decomposing diverse organic matters like phytoplankton-derived organic matters and chitin (Klippel et al., 2011), which may assist in digesting feeds in the gut of larvae as similar as *Rhodobacteraceae* taxa presumed to do (Wang, Y. et al., 2020). They also play roles in degrading excess feeds and shrimp excreta released in the rearing water, thus improving environmental quality of the culture system (Kim et al., 2022). Here, the hatching water largely shaped the *Flavobacteriaceae* assemblages of nauplii, and the contribution from the rearing water at the early stages was not as common as for the *Roseobacteraceae* assemblages; the role of internal succession in the later stages after *zoea* III was dominant but not as overwhelming as for the *Roseobacteraceae* assemblages. Furthermore, compared with *Roseobacteraceae*, more between-shrimp-strain differences in source composition of *Flavobacteriaceae* assemblages was observed. For example, frozen *Artemia* made a contribution for SR strain at *zoea* II, *mysis* (9 dph), *postlarvae* I, and *postlarvae* (13 dph) but only mattered at *zoea* II for FG strain. Collectively, these findings confirm taxonomic dependence and shrimp-strain specificity in assembling bacteria from complex sources into larval microbiota.

The monitoring and control of vibrios, as the most recognized bacterial pathogens in aquaculture, is critical in ensuring microbial safety across the entire life cycle of shrimps (Munkongwongsiri et al., 2022; Restrepo et al., 2021). Understanding the sources from which vibrios in the larval bacterial community are introduced is important for pathogen prevention and disease control. Here, the initial point of the rapid colonization of *Vibrionaceae* taxa from the rearing water to larval shrimps occurred at *nauplius* I and lasted until *zoea* I. The ubiquity of *Vibrionaceae* taxa in the rearing water during the mouth-opening stage of larvae indicates potential risk from opportunistic pathogens, though the taxonomic resolution the V4 region of the 16S rRNA gene could not meet the need of tracking specific pathogenic taxa down, which is worthy of future investigations by sequencing the 16S rRNA gene in full-length or metagenomics. After the mouth opening, frozen *Artemia* generally became the second important known source of *Vibrionaceae* taxa after internal succession, highlighting the necessity of proper treatment of *Artemia* to reduce pathogenic risk before feeding larval shrimp. The concern about microbial safety of using *Artemia* for larval feed in aquaculture has been raised and disinfection of *Artemia* before feeding has been proposed (Sorgeloos et al., 2001). Also, some practitioner used probiotics to precolonize *Artemia* and thus to prevent the propagation of pathogenic bacteria (Jamali et al., 2015). Our observation provides first micro-ecological evidence for cautious use of *Artemia* as an essential larval feed in aquaculture practices. It is generally believed that exchanging water with low toxic nitrogen concertation and bacterial density can help improve rearing water quality, but its potential in introducing pathogenic risk is often overlooked. Here, we found that the bacterial community in exchange water showed clear batch differences and contributed to larval *Vibrionaceae* assemblages at *postlarvae* (13 dph). Surprisingly, *Vibrionaceae* taxa from nauplii (*nauplius* I) continued to make a sustained contribution until the later stages (except for FG strain at *nauplius* VI - *zoea* I and *zoea* I, demonstrating that early life colonization of pathogenic bacteria may has a long-term impact on larval growth and development.

### 4.3 Determinism-dominated bacterial assembly from the sources into the larval bacterial community suggest strong host selection or screening

Our previous work has demonstrated that the assembly of larval bacterial community was overall more governed by neutral processes, including dispersal and ecological drift, when assuming larval bacterial metacommunity at each stage as the species pool for model inference (Wang, Y. et al., 2020). The main reason why aquatic animal diseases are difficult to control is the diversity of microbial sources and their complex transmission routes (Desrina et al., 2022; Rajeev et al., 2020). However, how are the bacteria from multiple sources assembled into the larval bacterial community cannot be distinguished by the previous assumption based on larval bacterial metacommunity. Here, we did source-tracking-oriented neutral model inferences and found that bacteria from previous-stage larvae tended to be deterministically assembled into the larval bacterial community at all the stages (based on the cumulative abundance of non-neutrally distributed ZOTUs vs. neutrally distributed ones). This suggests the portion of internally succeed bacteria likely experienced strong host selection or at least screening since the very beginning, even though the entire larval bacterial community was more governed by stochastic processes (Wang, Y. et al., 2020), which dominated bacterioplankton assembly in the rearing water as well (Heyse et al., 2021).

As similar as the above pattern observed in internal succession of bacteria by the host, bacteria in either environmental or input sources were also assembled into the larval bacterial community mainly by deterministic processes, regardless of shrimp strains. Determinism driven by host selection from the rearing water was previously found in most of the developmental stages of shrimp larvae (Wang, Y. et al., 2020). At the same time, given the overall minor contribution of external sources compared with internal succession after larval mouth opening, it seems that larval shrimps are very picky in choosing bacterial colonizer from any external sources here. Collectively, determinism-dominated assembly of bacteria from the sources into the larval bacterial community suggest strong host selection or screening, indicating that the regulation of host microbiota during larviculture could be achieved by enhancing deterministic assembly of beneficial taxa. Although some regulatory measures, like increasing C/N ratio by adding carbohydrate, to enhance assembly of beneficial bacterial groups such as *Rhodobacteraceae* taxa and thus maintaining a healthy bacterial community, have been proved effective in shrimp culture system (Guo et al., 2022), how to enhance deterministic assembly of targeted taxa in the larviculture system remains an open question.

After knowing that bacteria from either internal succession or external sources were overall assembled into the larval bacterial community *via* deterministic processes regardless of shrimp strains, revealing taxonomic preference of host selection from distinct sources and assessing commonness and difference between the two shrimp strains (with different culture traits and survival rates) in this aspect is critical for providing further insights into shrimp-strain-specific patterns in early life microbiota establishment (and potential microbial regulation strategies to improve survival rate of larval shrimps). As discussed above, the source composition of larval bacterial community was overall consistent between the two shrimp strains. At the same time, the taxonomic composition of bacteria retained by larvae of both strains *via* internal succession, as the major source of larval bacterial community, were overall similar with subtle different in relatively less abundant taxa. But the two shrimp strains showed common differences in taxonomic preference when selecting external bacteria from the same source(s) across the developmental stages. Studying effects of host genetics on the gut microbiome has been considered as a key to reveal how is gut microbiome involved in its host’s metabolism, immunity, and health (Sanna et al., 2022), yet only a few genetic loci have been consistently confirmed across multiple studies even in human. Therefore, how does genetic background influence host selection of microbiota and thus shaping larval shrimp microbiome is unknown. As we observed, the larvae of both strains tended to succeed a similar core microbiome mainly consisting of *Roseobacteraceae*, *Vibrionaceae*, *Flavobacteriaceae*, and *Halothiobacillaceae* after the mouth opening, which could be fundamental in maintaining homeostasis in the digestive tract, while larvae’s coping with the dynamics of environmental conditions and the transition of foods may result in shrimp-strain-specific selection of bacteria from external sources to meet their respective physiological needs due to distinct culture traits (phenotypes), which might also be related to the differentiation of survival rate between shrimp strains. Collectively, our study provides an insight into the potential impact of genetic background on host selection for bacteria during larviculture, proposing an important direction for future research on shrimp microbiome.

## Conclusions

To our best knowledge, this is the first systematic report on bacterial sources of *P. vannamei* larvae across the complete developmental cycle in a realistic larviculture practice. We found that the mouth opening is a turning point in bacterial source composition of larvae, and the large contribution from the rearing water at the beginning of mouth opening and the enhancing internal succession afterwards suggests the criticality and persistence of initial establishment of larval microbiota. While the dominance of external sources at *zoea* II suggests the importance of ensuring microbial safety at this vital sub-stage when *zoea*-II syndrome often occurs. To sum up, we recommend that the best practice for microbial regulation of larvae should be conducted through the rearing water (as the best pathway) as approaching *zoea* and at *zoea* (especially *zoea* II, as the best timing). Source tracking of potential beneficial and pathogenic bacterial groups suggests regulatory timing and routes of *Roseobacteraceae* (as the most abundant family with potential probiotics) and *Artemia* as a major source of vibrios. Determinism-dominated assembly of bacteria from either internal or external sources into larval community suggests strong host selection. Moreover, similarity in taxonomic preference of internal succession and frequent differences in taxonomic selection from external sources between the two shrimp strains indicate potential impact of genetic background on host selection for bacteria. Our findings provide ecological insights that facilitate microbial management and pathogen prevention during larviculture.

## Acknowledgements

This work is funded by Municipal Key R&D Program of Ningbo, China (2022Z177), Municipal Science & Technology Program of Wenzhou, China (KN20210008), K.C. Wong Magna Fund in Ningbo University, and Cultivation Platform of Zhejiang Key Laboratory of Intestinal Micro-ecology, Zhejiang Academy of Agricultural Sciences. The authors extend their sincere gratitude to the anonymous reviewers for their insightful comments and constructive suggestions, which have significantly enhanced the quality of this manuscript.

## Human and animal rights

All animal experiments were carried out in accordance with the U.K. Animals (Scientific Procedures) Act, 1986, and associated guidelines, EU Directive 2010/63/EU for animal experiments.

## Data availability

The sequence data have been submitted to the Sequence Read Archive of NCBI under accession number PRJNA1188736 and will be accessible with the following link after the acceptance of the manuscript: https://www.ncbi.nlm.nih.gov/sra/PRJNA1188736. Editors and reviewers can access the submitted data through the reviewer link: https://dataview.ncbi.nlm.nih.gov/object/PRJNA1188736?reviewer=nkviq9ucold7l06dm426k48m4l.

## Author contributions

K.W., D.Z., J.Y., and C.C conceived and organized the study; H.C and J.Y performed the larviculture and collected the samples; H.C. and F.Z. performed the bench work. K.W. and F.Z strategized the data analysis; F.Z., H.C., and R.C analyzed the data; K.W. and F.Z. wrote the manuscript. All the authors reviewed and approved the manuscript.

## References

Angthong, P., Chaiyapechara, S., Rungrassamee, W., 2023. Shrimp microbiome and immune development in the early life stages. Dev. Comp. Immunol. 147, 104765. 10.1016/j.dci.2023.104765.

Apprill, A., McNally, S., Parsons, R., Weber, L., 2015. Minor revision to V4 region SSU rRNA 806R gene primer greatly increases detection of SAR11 bacterioplankton. Aquat. Microb. Ecol. 75 (2), 129–137. 10.3354/ame01753.

Backhed, F., Roswall, J., Peng, Y., Feng, Q., Jia, H., Kovatcheva-Datchary, P., Li, Y., Xia, Y., Xie, H., Zhong, H., Khan, M.T., Zhang, J., Li, J., Xiao, L., Al-Aama, J., Zhang, D., Lee, Y.S., Kotowska, D., Colding, C., Tremaroli, V., Yin, Y., Bergman, S., Xu, X., Madsen, L., Kristiansen, K., Dahlgren, J., Wang, J., 2015. Dynamics and stabilization of the human gut microbiome during the first year of life. Cell Host Microbe. 17 (5), 690–703. 10.1016/j.chom.2015.04.004.

Bernal, M.G., Marrero, R.M., Campa-Córdova, Á.I., Mazón-Suástegui, J.M., 2016. Probiotic effect of *Streptomyces* strains alone or in combination with *Bacillus* and *Lactobacillus* in juveniles of the white shrimp *Litopenaeus vannamei*. Aquacult. Int. 25 (2), 927–939. 10.1007/s10499-016-0085-y.

Bi, Y., Cox, M.S., Zhang, F., Suen, G., Zhang, N., Tu, Y., Diao, Q., 2019. Feeding modes shape the acquisition and structure of the initial gut microbiota in newborn lambs. Environ. Microbiol. 21 (7), 2333–2346. 10.1111/1462-2920.14614.

Burns, A.R., Stephens, W.Z., Stagaman, K., Wong, S., Rawls, J.F., Guillemin, K., Bohannan, B.J., 2016. Contribution of neutral processes to the assembly of gut microbial communities in the zebrafish over host development. ISME J. 10, 655–664. 10.1038/ismej.2015.142.

Callac, N., Boulo, V., Giraud, C., Beauvais, M., Ansquer, D., Ballan, V., Maillez, J.R., Wabete, N., Pham, D., 2022. Microbiota of the rearing water of *Penaeus stylirostris* larvae influenced by lagoon seawater and specific key microbial lineages of larval stage and survival. Microbiol. Spectr. 10 (6), e0424122. 10.1128/spectrum.04241-22.

Chen, H., Zhang, F., Yu, J., Chen, R., Zhang, D., Chen, C., Wang, K., 2025. Divergence patterns of bacterial communities between larviculture systems of two *Penaeus vannamei* strains with distinct culture traits. Aquaculture 606, 742572. 10.1016/j.aquaculture.2025.742572.

Dai, W., Ye, J., Xue, Q., Liu, S., Xu, H., Liu, M., Lin, Z., 2023. Changes in bacterial communities of Kumamoto oyster larvae during their early development and following *Vibrio* infection resulting in a mass mortality event. Mar. Biotechnol. 25, 30–44. 10.1007/s10126-022-10178-0.

De Schryver, P., Vadstein, O., 2014. Ecological theory as a foundation to control pathogenic invasion in aquaculture. ISME J. 8, 2360–2368. 10.1038/ismej.2014.84.

Desrina, Prayitno, S.B., Verdegem, M.C.J., Verreth, J.A.J., Vlak, J.M., 2022. White spot syndrome virus host range and impact on transmission. Rev. Aquacult. 14 (4), 1843–1860. 10.1111/raq.12676.

Dinno, A., 2014. Package ‘dunn.test’: Dunn’s Test of Multiple Comparisons Using Rank Sums. https://cran.r-project.org/web/packages/dunn.test/.

Dong, P.S., Guo, H.P., Huang, L., Zhang, D.M., Wang, K., 2023. Glucose addition improves the culture performance of Pacific white shrimp by regulating the assembly of Rhodobacteraceae taxa in gut bacterial community. Aquaculture 567, 739254. 10.1016/j.aquaculture.2023.739254.

Duan, Y., Zhong, G., Nan, Y., Yang, Y., Xiao, M., Li, H., 2024. Effects of Nitrite Stress on the Antioxidant, Immunity, Energy Metabolism, and Microbial Community Status in the Intestine of Litopenaeus vannamei. Antioxidants 13 (11). 10.3390/antiox13111318.

Edgar, R.C., 2016. UNOISE2: improved error-correction for Illumina 16S and ITS amplicon sequencing. bioRxiv. 10.1101/081257.

Edgar, R.C., 2022. Muscle5: high-accuracy alignment ensembles enable unbiased assessments of sequence homology and phylogeny. Nat. Commun. 13, 6968. 10.1038/s41467-022-34630-w.

El-Saadony, M.T., Shehata, A.M., Alagawany, M., Abdel-Moneim, A.M.E., Selim, D.A., Abdo, M., Khafaga, A.F., El-Tarabily, K.A., El-Shall, N.A., Abd El-Hack, M.E., 2022. A review of shrimp aquaculture and factors affecting the gut microbiome. Aquacult. Int. 30 (6), 2847–2869. 10.1007/s10499-022-00936-1.

Elzhov, T.V., Mullen, K.M., Spiess, A.-N., Maintainer, B.B., 2016. minpack.lm. In: R Interface to the Levenberg-Marquardt Nonlinear Least-Squares Algorithm Found in MINPACK, Plus Support for Bounds. https://cran.r-project.org/web/packages/minpack.lm/minpack.lm.

Emerenciano, M.G.C., Rombenso, A.N., Vieira, F.D.N., Martins, M.A., Coman, G.J., Truong, H.H., Noble, T.H., Simon, C.J., 2022. Intensification of penaeid shrimp culture: an applied review of advances in production systems, nutrition and breeding. Animals 12 (3). 10.3390/ani12030236.

Fan, J., Chen, L., Mai, G., Zhang, H., Yang, J., Deng, D., Ma, Y., 2019. Dynamics of the gut microbiota in developmental stages of *Litopenaeus vannamei* reveal its association with body weight. Sci. Rep. 9, 734. 10.1038/s41598-018-37042-3.

FAO, 2024. Global aquaculture production 1950-2022. https://www.fao.org/fishery/statistics-query/en/aquaculture.

Gao, C.H., Chen, C., Akyol, T., Dusa, A., Yu, G., Cao, B., Cai, P., 2024. ggVennDiagram: intuitive venn diagram software extended. iMeta 3, e177. 10.1002/imt2.177.

Gundersen, M.S., Vadstein, O., De Schryver, P., Attramadal, K.J.K., 2022. Aquaculture rearing systems induce no legacy effects in Atlantic cod larvae or their rearing water bacterial communities. Sci. Rep. 12, 19812. 10.1038/s41598-022-24149-x.

Guo, H., Dong, P., Gao, F., Huang, L., Wang, S., Wang, R., Yan, M., Zhang, D., 2022. Sucrose addition directionally enhances bacterial community convergence and network stability of the shrimp culture system. npj Biofilms Microbi. 8, 22. 10.1038/s41522-022-00288-x.

Guo, H., Fu, X., He, J., Wang, R., Yan, M., Wang, J., Dong, P., Huang, L., Zhang, D., 2023. Gut bacterial consortium enriched in a biofloc system protects shrimp against *Vibrio parahaemolyticus* infection. Microbiome 11, 230. 10.1186/s40168-023-01663-2.

Hansen, G.H., Olafsen, J.A., 1999. Bacterial interactions in early life stages of marine cold water fish. Microb. Ecol. 38, 1–26. 10.1007/s002489900158.

He, Z., Pan, L., Zhang, M., Zhang, M., Huang, F., Gao, S., 2020. Metagenomic comparison of structure and function of microbial community between water, effluent and shrimp intestine of higher place *Litopenaeus vannamei* ponds. J. Appl. Microbiol. 129 (2), 243–255. 10.1111/jam.14610.

Healy, D.B., Ryan, C.A., Ross, R.P., Stanton, C., Dempsey, E.M., 2022. Clinical implications of preterm infant gut microbiome development. Nat. Microbiol. 7, 22–33. 10.1038/s41564-021-01025-4.

Heyse, J., Props, R., Kongnuan, P., De Schryver, P., Rombaut, G., Defoirdt, T., Boon, N., 2021. Rearing water microbiomes in white leg shrimp (*Litopenaeus vannamei*) larviculture assemble stochastically and are influenced by the microbiomes of live feed products. Environ. Microbiol. 23 (1), 281–298. 10.1111/1462-2920.15310.

Holt, C.C., Bass, D., Stentiford, G.D., van der Giezen, M., 2021. Understanding the role of the shrimp gut microbiome in health and disease. J. Invertebr. Pathol. 186, 107387. 10.1016/j.jip.2020.107387.

Huang, F., Pan, L., Song, M., Tian, C., Gao, S., 2018. Microbiota assemblages of water, sediment, and intestine and their associations with environmental factors and shrimp physiological health. Appl. Microbiol. Biotechnol. 102 (19), 8585–8598. 10.1007/s00253-018-9229-5.

Huang, L., Guo, H.P., Chen, C., Huang, X.L., Chen, W., Bao, F.J., Liu, W., Wang, S.P., Zhang, D.M., 2020. The bacteria from large-sized bioflocs are more associated with the shrimp gut microbiota in culture system. Aquaculture 523, 735159. 10.1016/j.aquaculture.2020.735159.

Hughes, K.R., Schofield, Z., Dalby, M.J., Caim, S., Chalklen, L., Bernuzzi, F., Alcon-Giner, C., Le Gall, G., Watson, A.J.M., Hall, L.J., 2020. The early life microbiota protects neonatal mice from pathological small intestinal epithelial cell shedding. FASEB J. 34 (5), 7075–7088. 10.1096/fj.202000042R.

Jamali, H., Imani, A., Abdollahi, D., Roozbehfar, R., Isari, A., 2015. Use of Probiotic *Bacillus* spp. in Rotifer (*Brachionus plicatilis*) and Artemia (*Artemia urmiana*) Enrichment: Effects on Growth and Survival of Pacific White Shrimp, *Litopenaeus vannamei*, Larvae. Probiotics Antimicrob. Proteins 7 (2), 118–125. 10.1007/s12602-015-9189-3.

Kembel, S.W., Cowan, P.D., Helmus, M.R., Cornwell, W.K., Morlon, H., Ackerly, D.D., Blomberg, S.P., Webb, C.O., 2010. Picante: R tools for integrating phylogenies and ecology. Bioinformatics 26 (11), 1463–1464. 10.1093/bioinformatics/btq166.

Kerr, C.A., Grice, D.M., Tran, C.D., Bauer, D.C., Li, D., Hendry, P., Hannan, G.N., 2015. Early life events influence whole-of-life metabolic health via gut microflora and gut permeability. Crit. Rev. Microbiol. 41 (3), 326–340. 10.3109/1040841X.2013.837863.

Kim, S.K., Song, J., Rajeev, M., Kim, S.K., Kang, I., Jang, I.K., Cho, J.C., 2022. Exploring bacterioplankton communities and their temporal dynamics in the rearing water of a biofloc-based shrimp (*Litopenaeus vannamei*) aquaculture system. Front. Microbiol. 13, 995699. 10.3389/fmicb.2022.995699.

Klippel, B., Lochner, A., Bruce, D.C., Davenport, K.W., Detter, C., Goodwin, L.A., Han, J., Han, S., Hauser, L., Land, M.L., Nolan, M., Ovchinnikova, G., Pennacchio, L., Pitluck, S., Tapia, R., Woyke, T., Wiebusch, S., Basner, A., Abe, F., Horikoshi, K., Keller, M., Antranikian, G., 2011. Complete genome sequences of *Krokinobacter* sp. strain 4H-3-7-5 and *Lacinutrix* sp. strain 5H-3-7-4, polysaccharide-degrading members of the family *Flavobacteriaceae*. J. Bacteriol. 193 (17), 4545–4546. 10.1128/JB.05518-11.

Knights, D., Kuczynski, J., Charlson, E.S., Zaneveld, J., Mozer, M.C., Collman, R.G., Bushman, F.D., Knight, R., Kelley, S.T., 2011. Bayesian community-wide culture-independent microbial source tracking. Nat. Methods 8 (9), 761–763. 10.1038/nmeth.1650.

Kronman, M.P., Zaoutis, T.E., Haynes, K., Feng, R., Coffin, S.E., 2012. Antibiotic exposure and IBD development among children: a population-based cohort study. Pediatrics 130 (4), e794–803. 10.1542/peds.2011-3886.

Kumar, R., Ng, T.H., Wang, H.C., 2020. Acute hepatopancreatic necrosis disease in penaeid shrimp. Rev. Aquacult. 12 (3), 1867–1880. 10.1111/raq.12414.

Kumar, T.S., Vidya, R., Alavandi, S.V., Vijayan, K.K., 2017. Zoea-2 syndrome of *Penaeus vannamei* in shrimp hatcheries. Aquaculture 479, 759–767. 10.1016/j.aquaculture.2017.07.022.

Liu, J.J., Wang, K., Wang, Y.T., Chen, W., Jin, Z.W., Yao, Z.Y., Zhang, D.M., 2019. Strain-specific changes in the gut microbiota profiles of the white shrimp in response to cold stress. Aquaculture 503, 357–366. 10.1016/j.aquaculture.2019.01.026.

Ma, T., Villot, C., Renaud, D., Skidmore, A., Chevaux, E., Steele, M., Guan, L.L., 2020. Linking perturbations to temporal changes in diversity, stability, and compositions of neonatal calf gut microbiota: prediction of diarrhea. ISME J 14 (9), 2223–2235. 10.1038/s41396-020-0678-3.

McKnight, D.T., Huerlimann, R., Bower, D.S., Schwarzkopf, L., Alford, R.A., Zenger, K.R., Jarman, S., 2018. Methods for normalizing microbiome data: an ecological perspective. Methods Ecol. Evol. 10 (3), 389–400. 10.1111/2041-210x.13115.

Munkongwongsiri, N., Prachumwat, A., Eamsaard, W., Lertsiri, K., Flegel, T.W., Stentiford, G.D., Sritunyalucksana, K., 2022. *Propionigenium* and *Vibrio* species identified as possible component causes of shrimp white feces syndrome (WFS) associated with the microsporidian *Enterocytozoon hepatopenaei*. J. Invertebr. Pathol. 192, 107784. 10.1016/j.jip.2022.107784.

O’Leary, N.A., Wright, M.W., Brister, J.R., Ciufo, S., Haddad, D., McVeigh, R., Rajput, B., Robbertse, B., Smith-White, B., Ako-Adjei, D., Astashyn, A., Badretdin, A., Bao, Y., Blinkova, O., Brover, V., Chetvernin, V., Choi, J., Cox, E., Ermolaeva, O., Farrell, C.M., Goldfarb, T., Gupta, T., Haft, D., Hatcher, E., Hlavina, W., Joardar, V.S., Kodali, V.K., Li, W., Maglott, D., Masterson, P., McGarvey, K.M., Murphy, M.R., O’Neill, K., Pujar, S., Rangwala, S.H., Rausch, D., Riddick, L.D., Schoch, C., Shkeda, A., Storz, S.S., Sun, H., Thibaud-Nissen, F., Tolstoy, I., Tully, R.E., Vatsan, A.R., Wallin, C., Webb, D., Wu, W., Landrum, M.J., Kimchi, A., Tatusova, T., DiCuccio, M., Kitts, P., Murphy, T.D., Pruitt, K.D., 2016. Reference sequence (RefSeq) database at NCBI: current status, taxonomic expansion, and functional annotation. Nucleic Acids Res 44 (D1), D733-D745. 10.1093/nar/gkv1189.

Oksanen, J., Blanche, F.G., Friendly, M., Kindt, R., Legendre, P., McGlinn, D., Minchin, P.R., O’Hara, R.B., Simpson, G.L., Solymos, P., Stevens, M., Szoecs, E., Wagner, H., 2013. vegan: community ecology package. https://cran.r-project.org/web/packages/vegan/.

Parada, A.E., Needham, D.M., Fuhrman, J.A., 2016. Every base matters: assessing small subunit rRNA primers for marine microbiomes with mock communities, time series and global field samples. Environ. Microbiol. 18 (5), 1403–1414. 10.1111/1462-2920.13023.

Pérez-Morales, A., Band-Schmidt, C.J., Martínez-Díaz, S.F., 2017. Mortality on zoea stage of the Pacific white shrimp *Litopenaeus vannamei* caused by *Cochlodinium polykrikoides* (Dinophyceae) and *Chattonella* spp. (Raphidophyceae). Mar. Biol. 164 (3), 57. 10.1007/s00227-017-3083-3.

Price, M.N., Dehal, P.S., Arkin, A.P., 2010. FastTree 2-approximately maximum-likelihood trees for large alignments. PLoS One 5 (3), e9490. 10.1371/journal.pone.0009490.

Racotta, I.S., Palacios, E., Herna n dez-Herrera, R., Bonilla, A., Pe rez-Rostro, C.I., ı rez, J.L.R., 2004. Criteria for assessing larval and postlarval quality of Pacific white shrimp (*Litopenaeus vannamei*, Boone, 1931). Aquaculture 233, 181-195. 10.1016/j.aquaculture.2003.09.031.

Racotta, I.S., Palacios, E., Ibarra, A.M., 2003. Shrimp larval quality in relation to broodstock condition. Aquaculture 227, 107–130. 10.1016/s0044-8486(03)00498-8.

Rajeev, M., Jung, I., Kang, I., Cho, J.C., 2024. Genome-centric metagenomics provides insights into the core microbial community and functional profiles of biofloc aquaculture. mSystems 9 (10), e0078224. 10.1128/msystems.00782-24.

Rajeev, R., Adithya, K.K., Kiran, G.S., Selvin, J., 2020. Healthy microbiome: a key to successful and sustainable shrimp aquaculture. Rev. Aquacult. 13, 238–258. 10.1111/raq.12471.

Restrepo, L., Dominguez-Borbor, C., Bajana, L., Betancourt, I., Rodriguez, J., Bayot, B., Reyes, A., 2021. Microbial community characterization of shrimp survivors to AHPND challenge test treated with an effective shrimp probiotic (*Vibrio diabolicus*). Microbiome 9, 88. 10.1186/s40168-021-01043-8.

Reyes, G., Betancourt, I., Andrade, B., Panchana, F., Roman, R., Sorroza, L., Trujillo, L.E., Bayot, B., 2022. Microbiome of *Penaeus vannamei* larvae and potential biomarkers associated with high and low survival in shrimp hatchery tanks affected by acute hepatopancreatic necrosis disease. Front. Microbiol. 13, 838640. 10.3389/fmicb.2022.838640.

Rungrassamee, W., Klanchui, A., Maibunkaew, S., Chaiyapechara, S., Jiravanichpaisal, P., Karoonuthaisiri, N., 2014. Characterization of intestinal bacteria in wild and domesticated adult black tiger shrimp (*Penaeus monodon*). PLoS One 9 (3), e91853. 10.1371/journal.pone.0091853.

Sanna, S., Kurilshikov, A., van der Graaf, A., Fu, J., Zhernakova, A., 2022. Challenges and future directions for studying effects of host genetics on the gut microbiome. Nat. Genet. 54, 100–106. 10.1038/s41588-021-00983-z.

Sanudo-Wilhelmy, S.A., Gomez-Consarnau, L., Suffridge, C., Webb, E.A., 2014. The role of B vitamins in marine biogeochemistry. Ann. Rev. Mar. Sci. 6, 339–367. 10.1146/annurev-marine-120710-100912.

Sha, H., Lu, J., Chen, J., Xiong, J., 2022. A meta-analysis study of the robustness and universality of gut microbiota-shrimp diseases relationship. Environ. Microbiol. 24 (9), 3924–3938. 10.1111/1462-2920.16024.

Shenhav, L., Thompson, M., Joseph, T.A., Briscoe, L., Furman, O., Bogumil, D., Mizrahi, I., Pe’er, I., Halperin, E., 2019. FEAST: fast expectation-maximization for microbial source tracking. Nat. Methods 16 (7), 627–632. 10.1038/s41592-019-0431-x.

Sloan, W.T., Lunn, M., Woodcock, S., Head, I.M., Nee, S., Curtis, T.P., 2006. Quantifying the roles of immigration and chance in shaping prokaryote community structure. Environ. Microbiol. 8 (4), 732–740. 10.1111/j.1462-2920.2005.00956.x.

Sonnenschein, E.C., Nielsen, K.F., D’Alvise, P., Porsby, C.H., Melchiorsen, J., Heilmann, J., Kalatzis, P.G., Lopez-Perez, M., Bunk, B., Sproer, C., Middelboe, M., Gram, L., 2017. Global occurrence and heterogeneity of the *Roseobacter*-clade species *Ruegeria mobilis*. ISME J. 11, 569–583. 10.1038/ismej.2016.111.

Sorgeloos, P., Dhert, P., Candreva, P., 2001. Use of the brine shrimp, *Artemia* spp., in marine fish larviculture. Aquaculture 200, 147–159. 10.1016/s0044-8486(01)00698-6.

Wang, R., Guo, Z., Tang, Y., Kuang, J., Duan, Y., Lin, H., Jiang, S., Shu, H., Huang, J., 2020. Effects on development and microbial community of shrimp *Litopenaeus vannamei* larvae with probiotics treatment. AMB Express 10, 109. 10.1186/s13568-020-01041-3.

Wang, Y., Wang, K., Huang, L., Dong, P., Wang, S., Chen, H., Lu, Z., Hou, D., Zhang, D., 2020. Fine-scale succession patterns and assembly mechanisms of bacterial community of *Litopenaeus vannamei* larvae across the developmental cycle. Microbiome 8, 106. 10.1186/s40168-020-00879-w.

Xiong, J., Xuan, L., Yu, W., Zhu, J., Qiu, Q., Chen, J., 2019. Spatiotemporal successions of shrimp gut microbial colonization: high consistency despite distinct species pool. Environ. Microbiol. 21 (4), 1383–1394. 10.1111/1462-2920.14578.

Yan, H.Z., Lin, D.D., Gu, G.K., Huang, Y.J., Hu, X.Y., Yu, Z.H., Hou, D.D., Zhang, D., Campbell, B.J., Wang, K., 2024. Taxonomic dependency and spatial heterogeneity in assembly mechanisms of bacteria across complex coastal waters. Ecol. Process 13, 6. 10.1186/s13717-023-00480-7.

Zhang, X.C., Li, X.H., Lu, J.Q., Qiu, Q.F., Chen, J., Xiong, J.B., 2021. Quantifying the importance of external and internal sources to the gut microbiota in juvenile and adult shrimp. Aquaculture 531, 735910. 10.1016/j.aquaculture.2020.735910.

