## Supplemental information for "Source tracking of larval bacterial community of Pacific white shrimp across the developmental cycle: ecological insights for microbial management and pathogen prevention in larviculture"

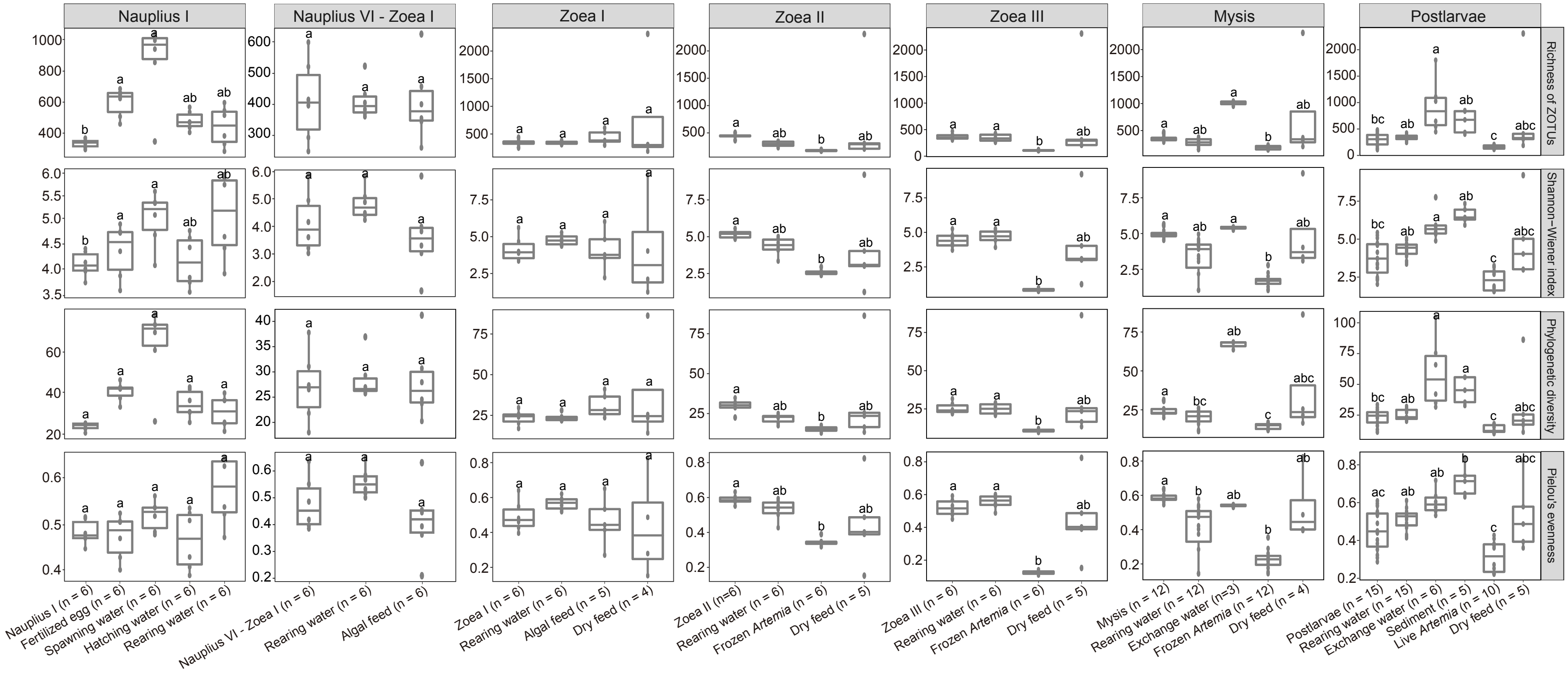


**Figure S1** Alpha-diversity and evenness indices of bacterial communities of shrimp larvae and their potential sources across host developmental stages. The boxes illustrate the median and interquartile range, and the whiskers extend from the minimum to the maximum values. Data with different letters indicate significant differences after multiple comparisons of Kruskal-Wallis test ($P < 0.05$).

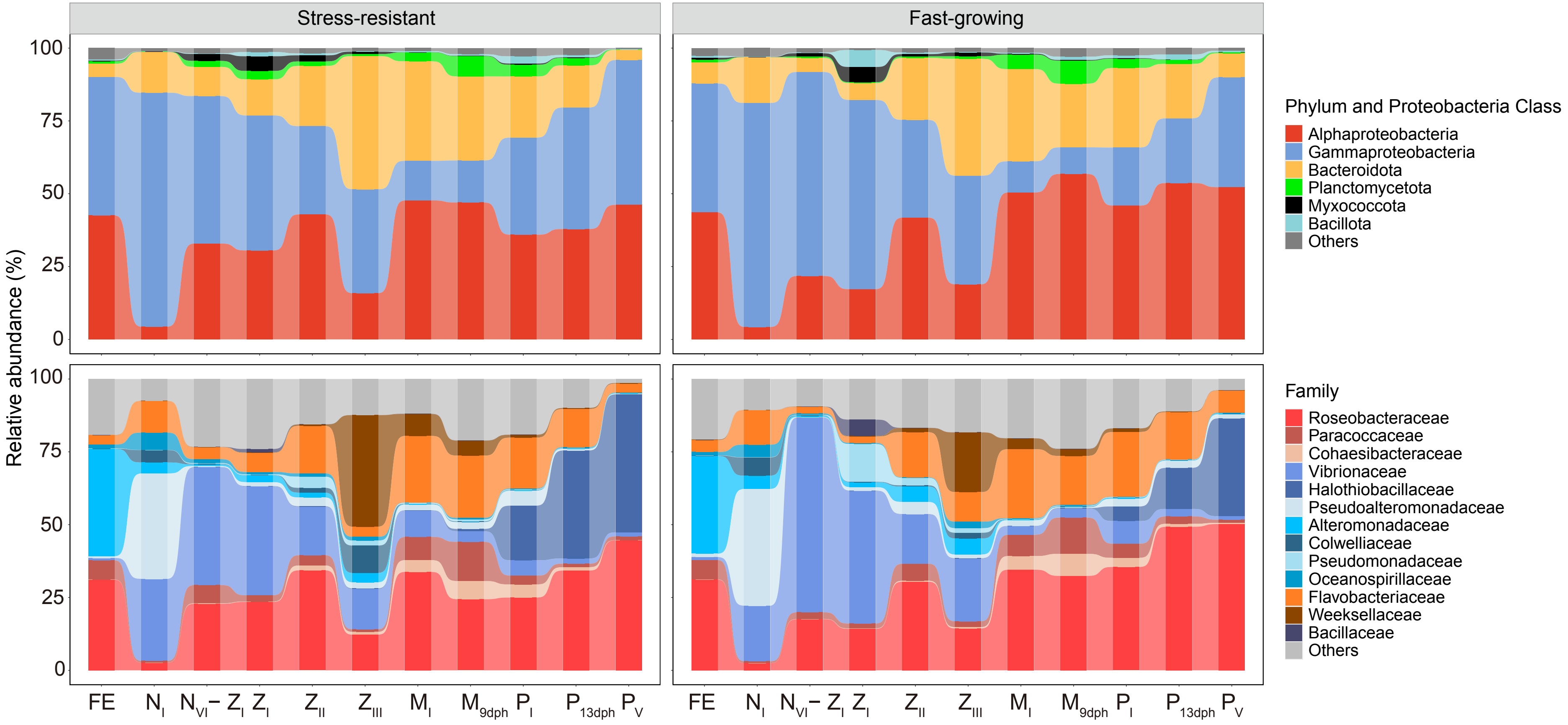


**Figure S2** Sankey diagram showing the dynamics of relative abundance of the dominant bacterial phyla and proteobacterial classes (average relative abundant > 5% at least in one sampling day (**upper**) and families (average relative abundant > 5% at least in one sampling day (**Bottom**) fertilized egg and larvae over host development. FE, fertilized egg; $N_I$, *nauplius* I; $N_{VI}$-$Z_I$, *nauplius* VI - *zoea* I; $Z_I$, *zoea* I; $Z_{II}$, *zoea* II, $Z_{III}$, *zoea* III; $M_I$, *mysis* I; $M_{9dph}$, *mysis* (9 dph); $P_I$, *postlarvae* I; $P_{13dph}$, *postlarvae* (13 dph); $P_V$, *postlarvae* V.

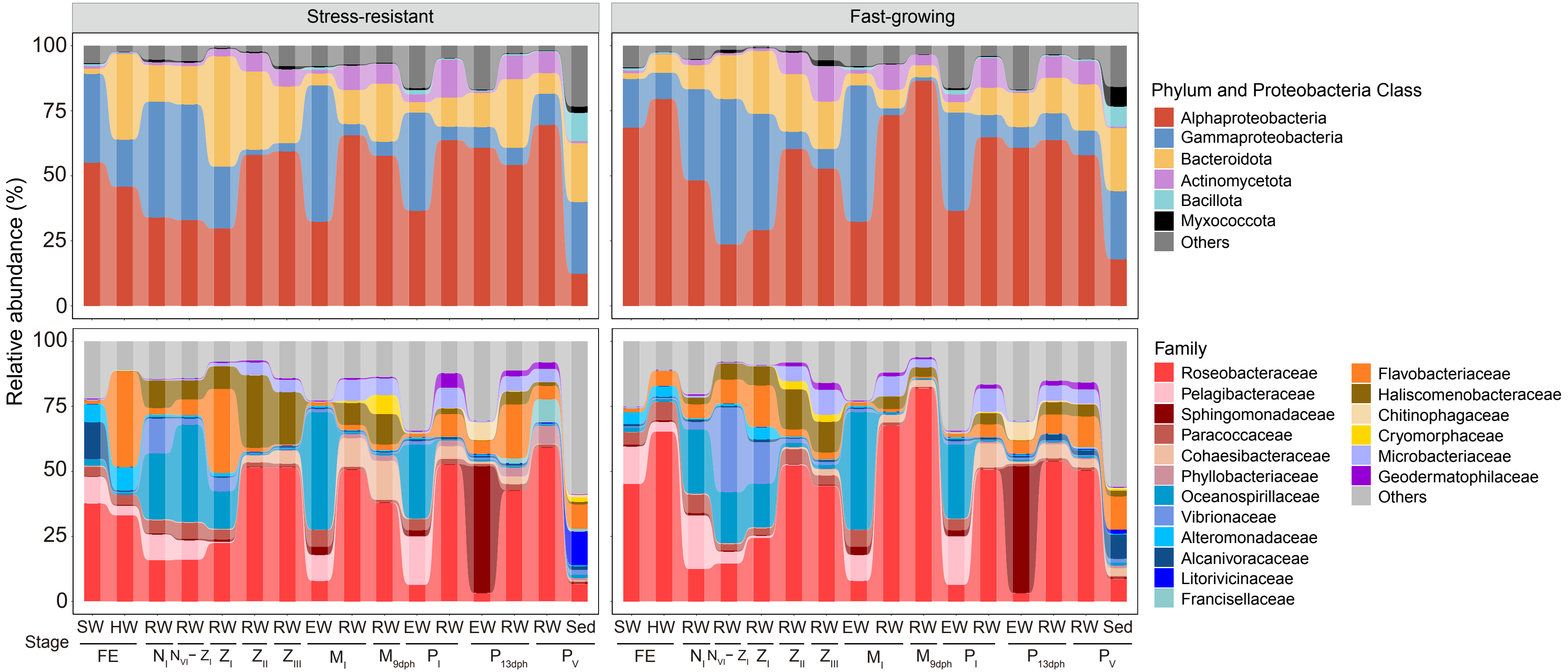


**Figure S3** Sankey diagram showing the dynamics of relative abundances of the dominant bacterial phyla and proteobacterial classes (average relative abundant > 5% at least in one sampling day (**Upper**) and families (average relative abundant > 5% at least in one sampling day (**Bottom**) in environmental sources including spawning water (SW), hatching water (HW), rearing water (RW), exchange water (EW), and sediment (Sed) across host developmental stages. FE, fertilized egg; $N_I$, *nauplius* I; $N_{VI}$-$Z_I$, *nauplius* VI - *zoea* I; $Z_I$, *zoea* I; $Z_{II}$, *zoea* II, $Z_{III}$, *zoea* III; $M_I$, *mysis* I; $M_{9dph}$, *mysis* (9 dph); $P_I$, *postlarvae* I; $P_{13dph}$, *postlarvae* (13 dph); $P_V$, *postlarvae* V.

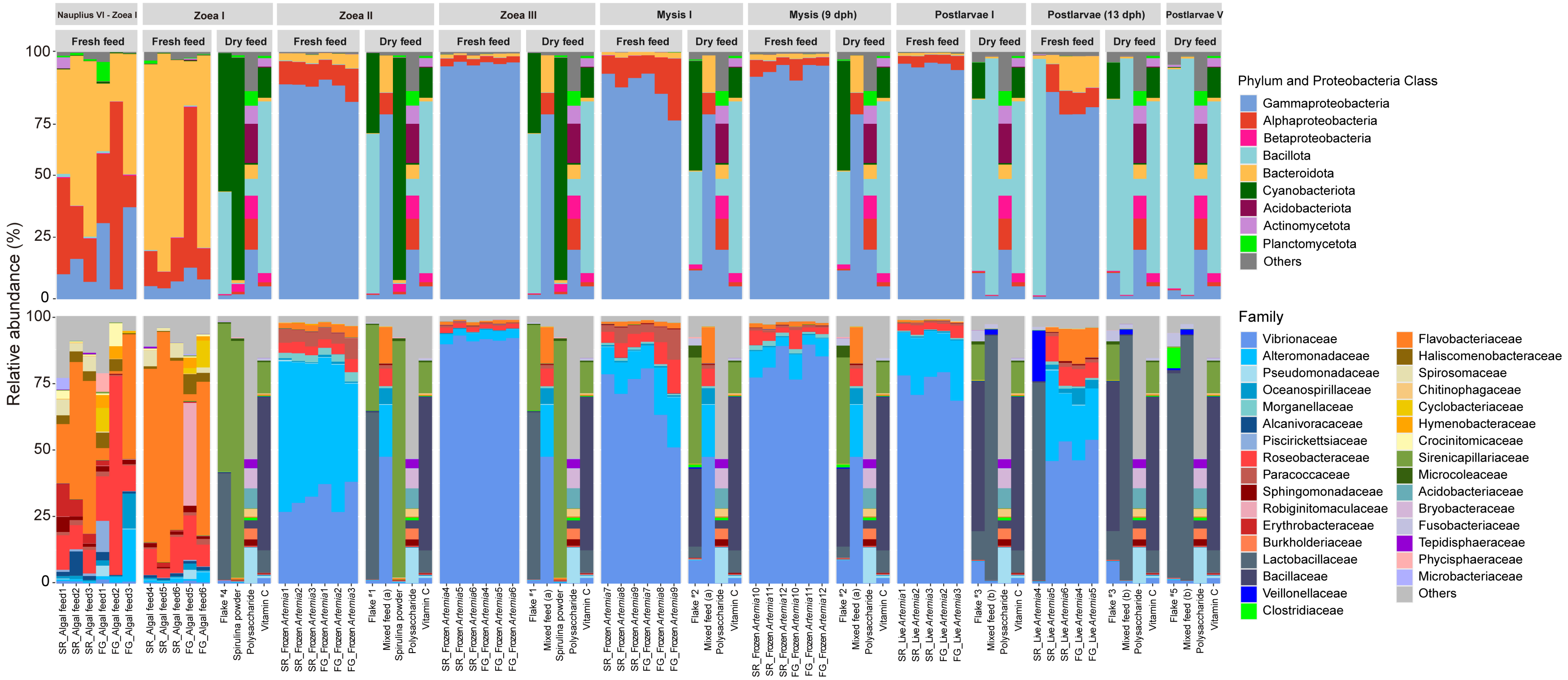


**Figure S4** Relative abundance of the dominant bacterial phyla and proteobacterial classes (average relative abundant > 2% at least in one sampling day) (**Upper**) and families (average relative abundant > 2% at least in one sampling day) (**Bottom**) in input sources, including fresh feeds (algal feed, frozen *Artemia*, and live *Artemia*) and dry feeds (flakes #1–#5, mixed feed (types a and b), spirulina powder, polysaccharide, and vitamin C) across host developmental stages. For each type of dry feed, a single representative sample—sourced from the same package as all others of its type—was used for DNA extraction and sequencing, and served as the reference across all sampling stages for comparative purposes. SR, stress-resistant shrimp strain; FG, fast-growing shrimp strain.

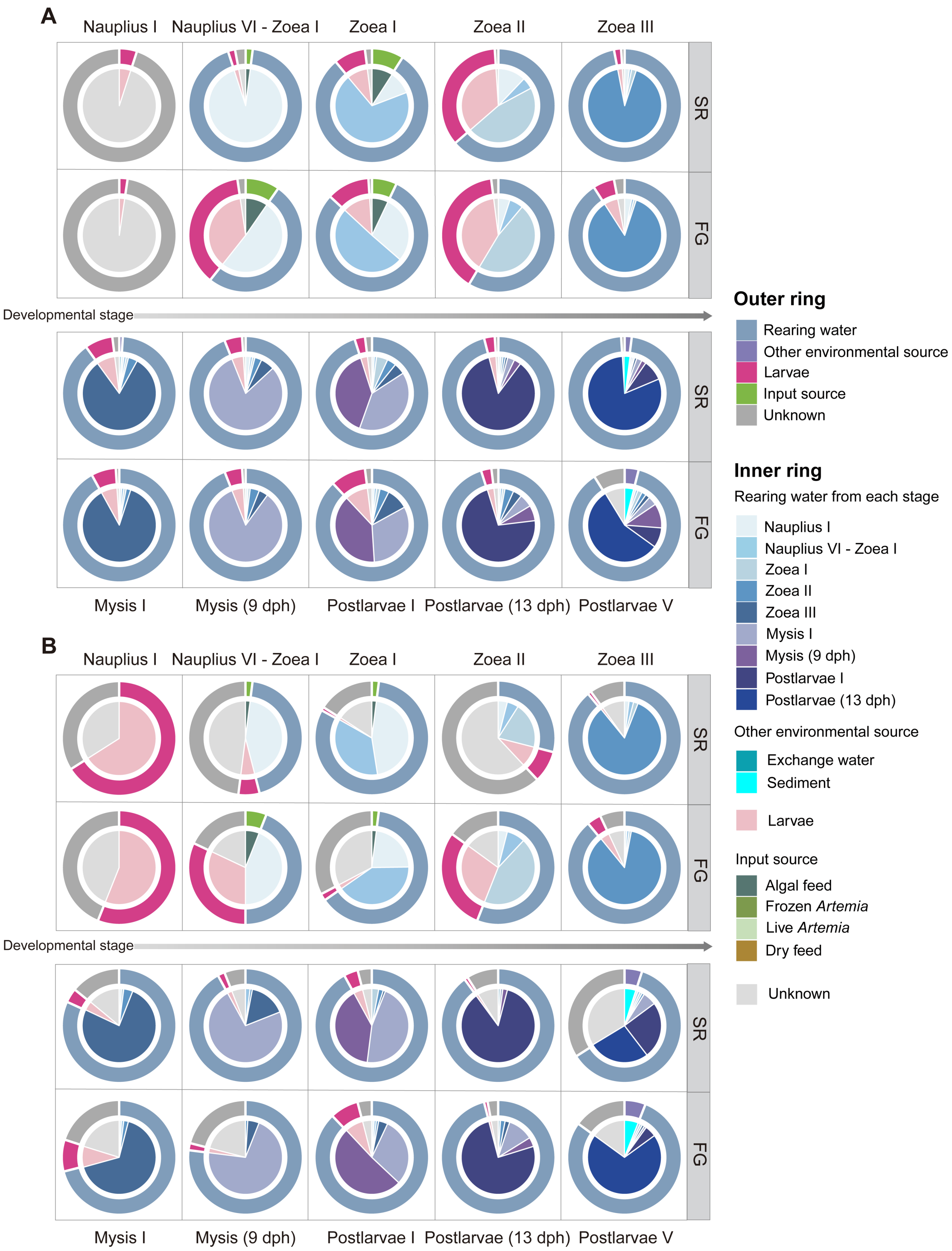


**Figure S5** The relative contribution of various potential sources to the bacterial community of rearing waters across host developmental stages inferred by SourceTracker (A) and Fast Expectation-maximization for Microbial Source Tracking (FEAST; B). Larvae present larval samples of a given stage corresponding to the rearing water. SR, stress-resistant shrimp strain; FG, fast-growing shrimp strain.
